# Generative inference unifies feedback processing for learning and perception in natural and artificial vision

**DOI:** 10.1101/2025.10.21.683535

**Authors:** Tahereh Toosi, Kenneth D. Miller

## Abstract

We understand how neurons respond selectively to patterns in visual input to support object recognition; however, how these circuits support perceptual grouping, illusory percepts, and imagination is not understood. These perceptual experiences are thought to require combining what we have learned about the world with incoming sensory signals, yet the neural mechanism for this integration remains unclear. Here we show that networks tuned for object recognition implicitly learn the distribution of their input, which can be accessed through feedback connections that tune synaptic weights. We introduce Generative Inference, a computational framework in which feedback pathways that adjust connection weights during learning are repurposed during perception to combine learned knowledge with sensory input, fulfilling flexible inference goals such as increasing confidence. Generative Inference enables networks tuned solely for recognition to spontaneously produce perceptual grouping, illusory contours, shape completion, and pattern formation resembling imagination, while preserving their recognition abilities. The framework reproduces neural signatures observed across perceptual experiments: delayed responses in feedback-receiving layers of early visual cortex that disappear when feedback connections are disrupted. We show that, under stated assumptions, gradients of classification error approximate directions that are informative about the data distribution, establishing a theoretical connection between recognition and generation. Together, these findings show that pattern recognition and pattern generation rely on a shared computational substrate through dual use of feedback pathways. This principle explains how neural systems recognize familiar objects reliably while remaining flexible enough to interpret incomplete or ambiguous information, and suggests that reusing learning signals for perception may be a general feature of both biological brains and artificial networks.

## Main

Artificial neural networks (ANNs) show remarkable convergence with both brain activation patterns across various domains. Neural activity patterns in visual cortex are accounted for by hierarchical vision models trained on object recognition^1–3^, neural activity patterns in auditory cortex are captured by spectro-temporal models trained on auditory tasks^4^, and neural activity patterns in language regions are explained by transformer models trained on language processing^5;6^. Despite this striking alignment between artificial and biological neural representations, a crucial mystery remains: the rich phenomenology of human perception bears no clear relationship to the computational operations of these models. Visual illusions and Gestalt principles reveal this explanatory gap most starkly: we perceive vivid contours, coherent objects, and meaningful structures that exist nowhere in the physical stimulus. What accounts for this failure of neural networks that otherwise successfully capture the information-processing machinery of vision?

The answer may lie in recognizing that while models excel at stimulus-driven pattern recognition, they lack mechanisms to integrate learned priors with incoming sensory evidence, a process central to perception. Human perception does not operate as a passive feature detector that simply registers whatever patterns stimulate the retina. Instead, it is an inference system that constructs interpretations by combining sensory data with prior knowledge. What we experience as distinct visual phenomena, from precise object recognition to the ethereal constructions of imagination, actually represent points along a single computational continuum (Figure 1 **A**). At one end, we find rapid, bottom-up processing that enables immediate recognition of clear visual signals^7^. At the other end lie the purely top-down constructions of mental imagery and dreams, where the visual system generates percepts independent of external input. The critical domain lies between these extremes: a rich middle ground where perception arises from a sophisticated inferential process that integrates sensory data with learned priors to fill gaps, resolve ambiguities, and extract coherent interpretations that the brain determines are implied by the sensory input.

**Figure 1:**
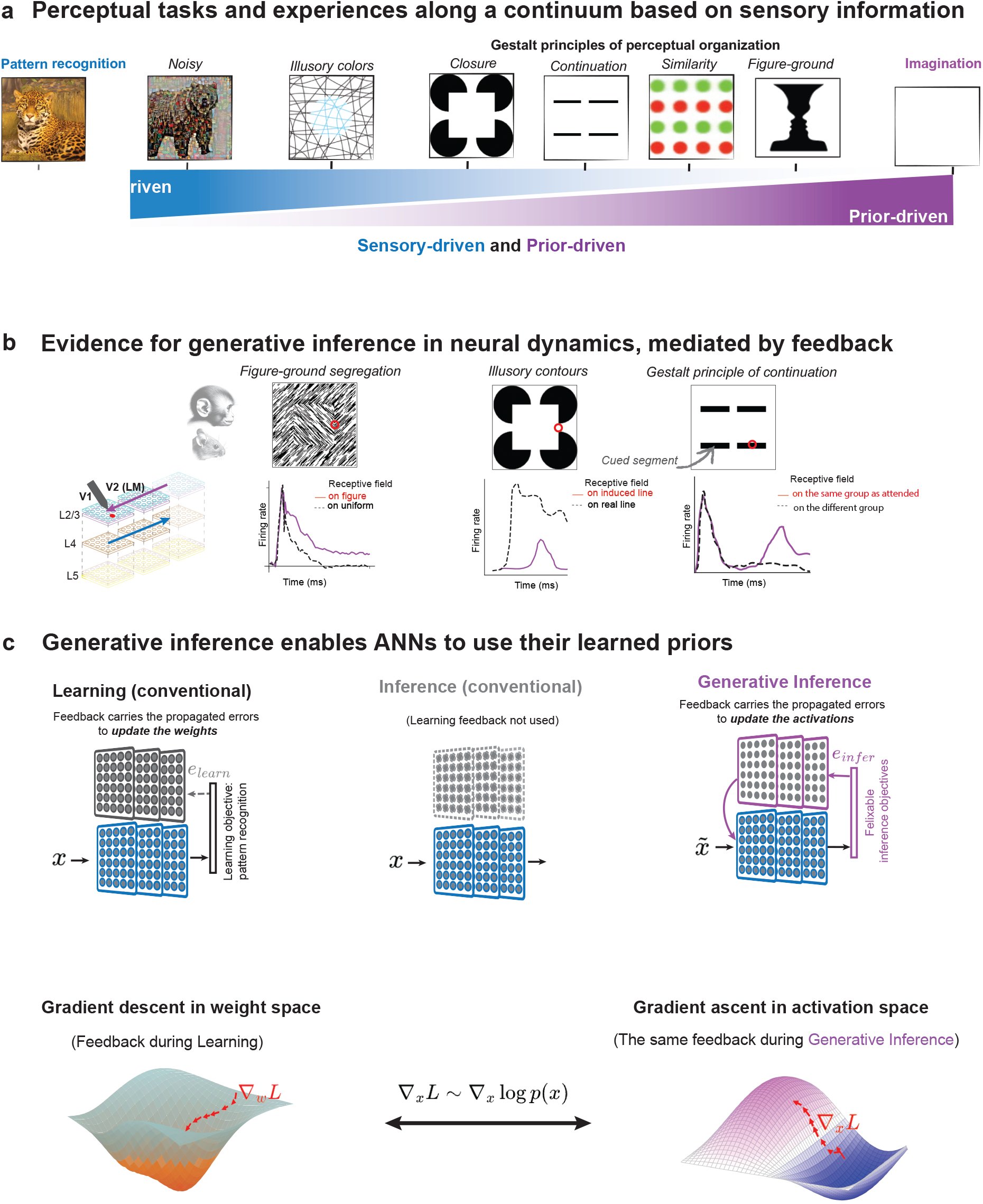
Generative Inference as a unifying principle for feedback processing. **a** Visual perception tasks and experiences can be located on a continuum defined by degree of reliance on sensory evidence. As sensory evidence decreases, the integration of priors becomes increasingly crucial. This spectrum unifies seemingly disparate visual phenomena: from fast core object recognition, demanding minimal prior integration^1;7^ through recognition despite noise or occlusion^10–12^, or instances of Gestalt principles of organization such as figure-ground segregation^17;25^, illusory contours^14–16;19^, colors, or shapes^9^, and ultimately imagination (entirely prior-dependent). While standard artificial neural networks excel at the sensory-dominated end of this spectrum, they typically fail at tasks requiring substantial prior integration^26;27^. **b** Neural dynamics reflecting prior integration reveal consistent feedback-mediated responses across diverse perceptual phenomena. Activity patterns in early visual cortices show characteristic laminar-specific signatures, with stronger modulation in feedback-receiving layers (L2/3) compared to input layers (L4). Similar feedback-driven temporal dynamics appear for figure-ground segregation^25^, illusory contours^15;16;19^, and Gestalt grouping principles of grouping^28^. **c** (*Top*) In conventional learning (left), feedback carries error signals *e*_*learn*_ to update synaptic weights^20^. During standard inference (middle), feedback connections which were previously used for learning are unused. Generative Inference (right) activates these same pathways to update neuronal activations based on iterative adjustments using errors for a flexible inference objective *e*_*infer*_, enabling dynamic integration of sensory evidence with learned priors. This repurposing of feedback pathways allows networks to access their learned statistical regularities during perception. (*Bottom*) During learning (left), feedback pathways carry gradients ∇_*W*_*L* for weight optimization via gradient descent. Leveraging the theoretical link we establish between gradients of loss with respect to input and gradients of the data distribution (∇_*x*_*L* ≈ ∇_*x*_ log *p*(*x*)), Generative Inference enables the same feedback pathways to carry data distribution gradients that guide activation optimization via gradient ascent (right). This allows neural activity to navigate toward regions of higher probability under the learned statistical priors (See Supplementary A).

Converging evidence suggests that the brain integrates learned priors with incoming sensory input through a generative process mediated by feedback from higher visual areas, especially when perception cannot rely on feedforward sensory cues alone. This evidence shows that, while clear, undistorted stimuli elicit rapid, primarily feedforward-driven responses^7;8^, ambiguous^9^, incomplete^10–12^ or unusual stimuli^13^ engage slower, feedback-dependent mechanisms that exhibit characteristic delayed responses and laminar-specific activation patterns^14–17^. These experiments reveal three hallmark signatures of the prior integration process (Figure 1**B**): (1) **temporal delays** of 70–100ms for ambiguous stimuli in early visual areas compared to 30–40ms for real contours^15;17^, (2) **feedback dependence**, where optogenetic inhibition of higher areas eliminates responses to illusory but not real stimuli^16;18^, and (3) **laminar specificity**, with responses concentrated in feedback-receiving layers (L2/3, L5/6) rather than the primary input layer (L4)^15;16;19^.

We hypothesize that feedback pathways serve dual computational roles in cortical processing. During learning, feedback carries error signals that guide synaptic modification^20^, while during perception, the same pathways can implement inference processes as discussed above. Here we introduce the computational principle of Generative Inference, which formalizes this dual role, demonstrating how the same feedback pathways used during learning can be repurposed for inference (Figure 1**C**). During learning, feedback carries error signals to update synaptic weights, whether through gradient descent, as used in the backpropagation algorithm in ANNs and in postulated biological implementations^21^, or through yet unknown biological algorithms. In current ANNs, these feedback connections remain unused during inference (which, in ANNs, means the process of producing network outputs with fixed weights, *i*.*e*. without learning; what in biology would simply be called perception). Generative Inference demonstrates that these same pathways can be activated during inference to iteratively update neural activations rather than weights, enabling dynamic integration of sensory evidence with learned priors. This reproduces a wide array of perceptual phenomena. While we demonstrate this in ANNs using backpropagation to drive feedback connections, we hypothesize that whatever algorithms are used to drive feedback-guided learning in biological systems can similarly be repurposed to integrate priors during perception.

This repurposing is theoretically grounded in the mathematical equivalence between gradient descent in weight space during learning and gradient ascent in activation space during inference (Supplementary A). During learning, feedback pathways carry gradients ∇_*W*_*L* that optimize weights via gradient descent. Our framework leverages the established theoretical link between gradients of classification loss and the score function of generative models (∇_*x*_*L* ≈ ∇_*x*_*logp*(*x*)) to show that these same feedback pathways can carry data distribution gradients during inference. This enables neural activity to navigate toward regions of higher probability under the learned statistical priors, effectively implementing score-based generative inference^22^ through the network’s existing architecture. To test this theoretical framework empirically, we examined whether Generative Inference can account for the diverse perceptual phenomena where feedback modulates neural activity in biological vision. We focus on key domains where feedforward/recurrent models fail but are hallmarks of biological neural networks: Gestalt principles of perceptual organization, illusory perception, figure-ground segregation, and imagination.

### Generative inference: a unified framework for feedback processing

We demonstrate the capabilities of Generative Inference using the classic Kanizsa square illusion^13^, in which four inducer segments positioned at the corners of an imaginary square evoke the perception of an illusory occluding shape. Neurophysiological studies have shown that neurons in early visual cortex respond to these non-existent contours, with activity emerging 70–100 ms after responses to real edges^15^, and localized to feedback-receiving layers^15;19^, suggesting a top-down generative process. To test whether this phenomenon can emerge in artificial neural networks, we present the Kanizsa square to a robustly trained convolutional network^23;24^ and apply our Generative Inference framework, reusing the same feedback pathways involved in training to perform inference-time updates. This process—Generative Inference—progressively refines the internal representation of the stimulus, producing activation patterns that reveal the presence of the illusory square over time (Figure 2A–B).

**Figure 2:**
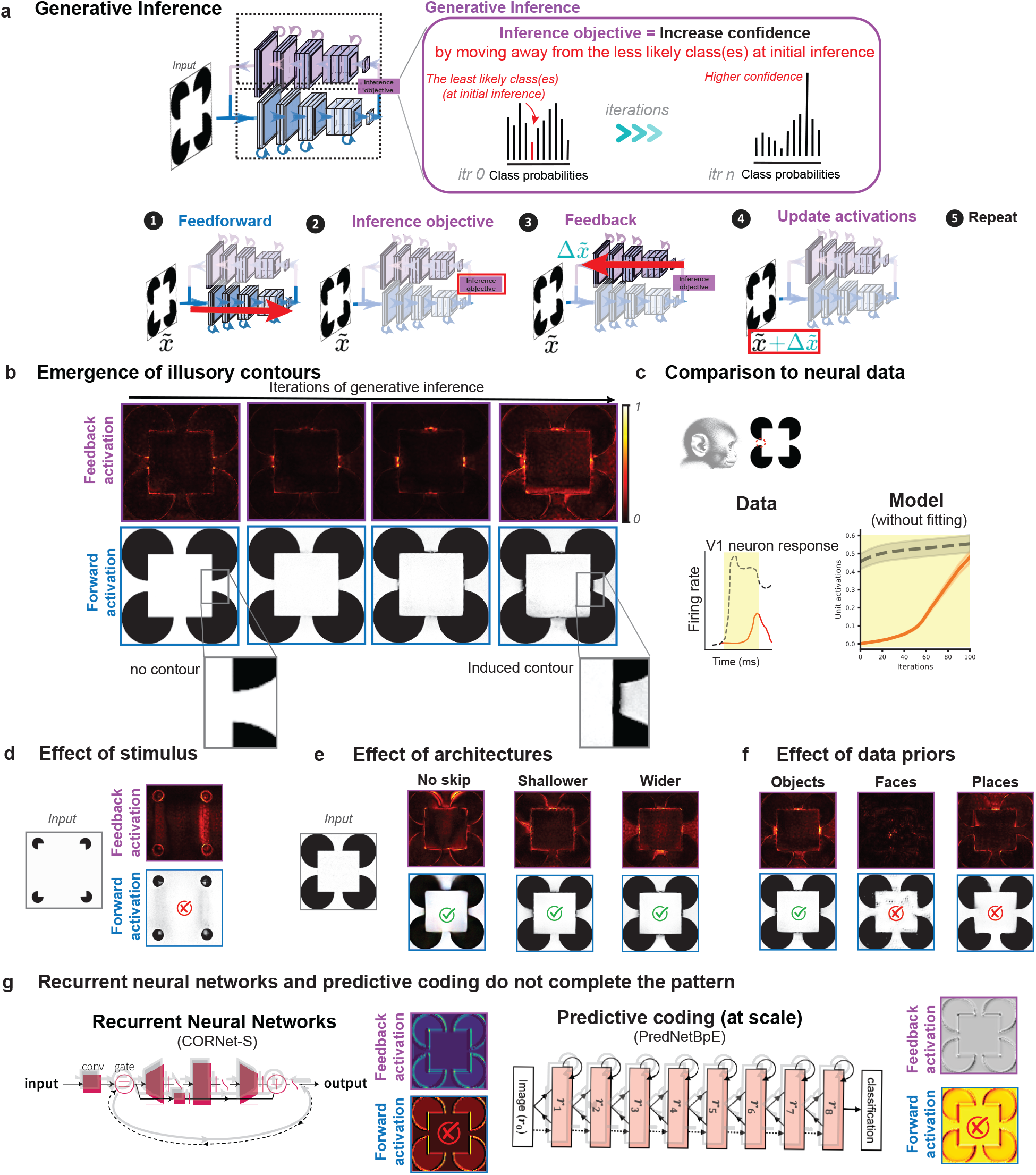
Generative Inference produces illusory contour perception through feedback processing. **a** Kanizsa square^13^ is presented to a generic convolutional neural network^23^ trained for robust object recognition^29^ on ImageNet^30^. Generative Inference reuses the feedback pathway involved in training to update activations at inference time, driven by a novel inference objective: increase confidence. Generative Inference implements an iterative procedure: (1) Feedforward pass: compute initial class probabilities and identify the least confident class; (2) Inference objective: select the least confident class as target for gradient computation; (3) Feedback error propagation: compute gradients of negative log-likelihood with respect to input activations, using the same feedback pathways employed during training; (4) Constrained activation update: move representations away from the least confident class by applying normalized gradients plus stochastic noise, while constraining updates within an *ϵ*-ball of the original input; (5) Iteration control: repeat steps 2-4 for *T* iterations. This objective iteratively shifts the network’s internal representations away from less likely classes—those initially assigned low confidence—toward more confident interpretations consistent with learned priors. Black dashed line shows the temporal profile of neural responses to real contours, while red line shows the delayed emergence of responses to illusory contours. **b** Progressive emergence of illusory contours through feedback processing. Top row shows feedback activation patterns evolving over iterations, while bottom row shows corresponding forward activations, demonstrating how the Kanizsa square gradually emerges.^14–16;19;31^. **c** Comparison between neural data and model results, showing how illusory contour responses emerge with a characteristic delay compared to responses to real contours, matching observations in biological visual systems. **d** Control experiment showing specificity: when inducer segments are placed too far apart, illusory contours fail to form, consistent with perceptual thresholds observed in humans^32^. **e** Different network architectures all demonstrate contour completion but with distinct activation patterns, showing the generality of the effect (Small, No skip connections: vgg-16, Shallower:resnet18, Wider:resnet50-Wide). **f** The effect of training data priors: networks with identical architecture (ResNet50) but trained on different datasets show varying abilities to generate illusory contours. Networks trained on Object recognition (ImageNet) readily generate the illusion, while those with the same architecture but trained only on faces ^33^ or scenes from places^34^ fail to do so, demonstrating that the appropriate statistical priors are essential for illusory contour perception. **g** Recurrent neural networks and predictive coding at scale (both large scale and trained on ImageNet) fail to complete the edges even though they benefit from feedback processing (see Supp. F).

**Figure 3:**
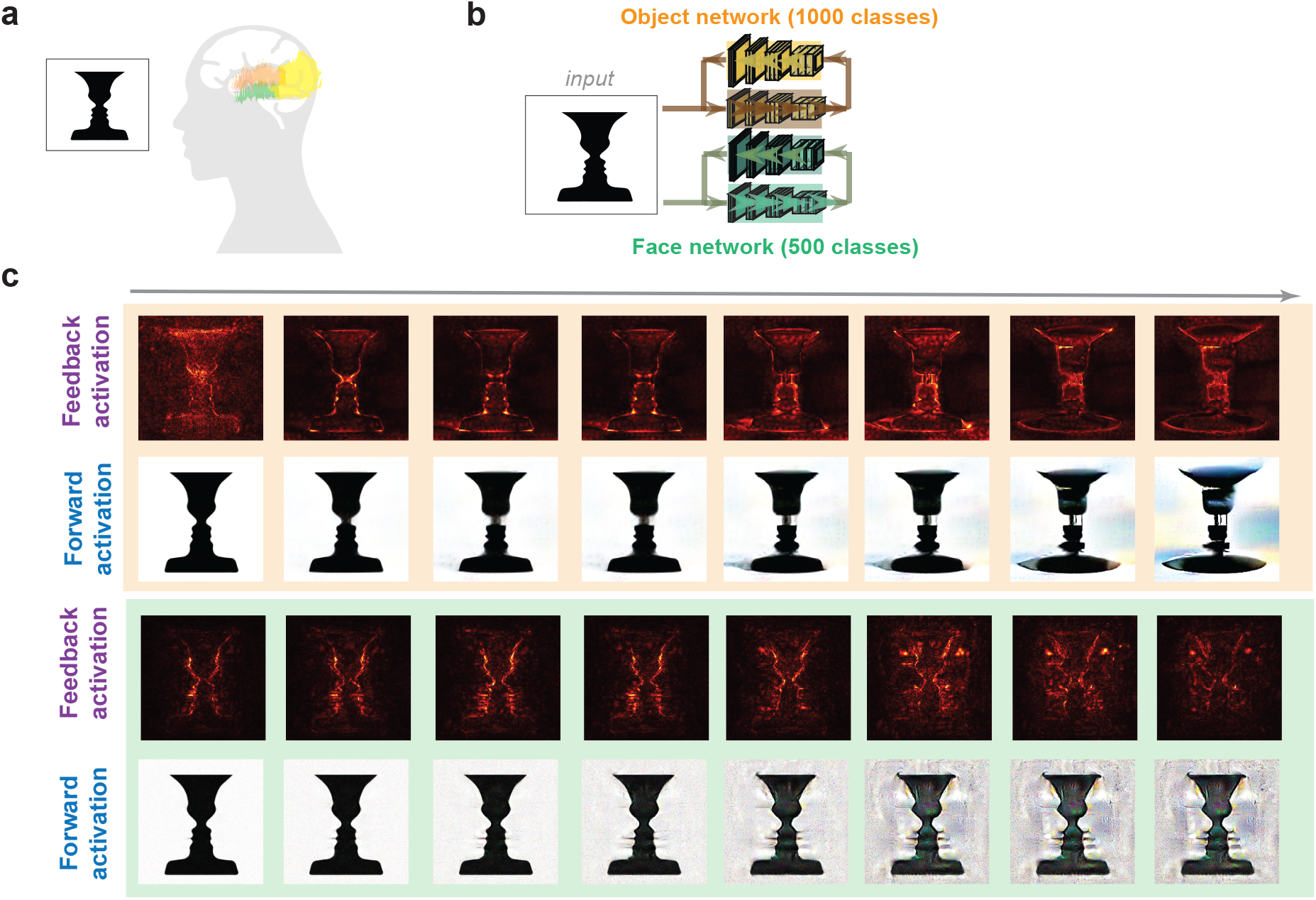
Bistable perception emerges from accumulating learned priors through feedback processing. **a** The Rubin face-vase illusion is an example of bistable perception, where the same visual stimulus can be interpreted as either two faces in profile or a central vase, with perception spontaneously alternating between these interpretations. Neuroimaging studies show that early visual brain areas (including primary visual cortex) reflect the observer’s current perceptual state rather than just the physical stimulus, with neural activity varying according to whether participants report seeing faces or the vase despite identical retinal input^9^. **b** We trained a ResNet50 network for face recognition (VGGFace2, 500 classes) and used it along an identical architecture trained on object recognition (ImageNet, 1000 classes). Critically, neither network was exposed to any instance of Rubin’s vase stimuli during training, ensuring that perceptual interpretations emerge purely from learned statistical priors. **C** When presented with an ambiguous face-vase stimulus, networks with different priors generate distinct interpretations through feedback. Top: A network trained for object recognition develops vase-like features. Bottom: A network, with the same architecture, ResNet50^23^, trained for face recognition generates face-like patterns. For each, evolving activation patterns are shown across feedback iterations (left to right).

Here, Generative inference uses an inference objective which is different from the learning objective: rather than optimizing weights for a fixed category label, the goal of inference here is to refine activations by moving away from the least likely class(es) at initial prediction. This “increase confidence” objective effectively pulls the network’s internal activations toward regions of representation space that are more consistent with the learned data distribution (Supplementary B.1). As feedback updates are applied over multiple iterations, activation patterns across layers become increasingly structured, and illusory features—absent during the initial feedforward pass—emerge naturally from the model’s own priors. This process provides a computational analogue of perceptual completion, mimicking the delayed and structured responses observed in the primate brain when perceiving illusory contours (Figure 2 C).

To validate the specificity and robustness of these effects, we conduct a series of control experiments. When the inducers are positioned too far apart, no illusory contour emerges, matching human vision (Figure 2D). We also show that the effect is architecture-agnostic: networks with differing structures (e.g., with or without skip connections, shallower or wider variants) still exhibit generative contour completion, although the resulting patterns differ (Figure 2E). Critically, the presence of the illusion depends on the statistical priors learned during training: models with identical architecture trained on different datasets (e.g. Faces or Places) fail to generate the Kanizsa square, whereas those trained on object-centric datasets (e.g., ImageNet) succeed (Figure 2F). These findings highlight that both structural stimulus cues and learned priors are essential for generating perceptual inferences, paralleling observations in biological systems and supporting the view that feedback-driven generative inference is a plausible mechanism for visual completion.

### Generative inference explains the interpretation of ambiguous inputs by iterative accumulation of priors

The Kanizsa square is often cited as a textbook example of the Gestalt principle of closure, where the brain perceptually completes missing edges to form a coherent shape. But closure is only one of several organizational principles Gestalt theory uses to explain perception. This raises a broader question: can a single computational mechanism, such as Generative Inference, also explain other perceptual phenomena traditionally attributed to Gestalt principles, including figure-ground segregation, grouping by similarity and proximity, and contextual illusions of brightness and color?

A particularly well-studied case of perceptual ambiguity is the Rubin face-vase illusion, where the same visual input can be interpreted either as two faces in profile or as a central vase. Functional imaging studies^9^ show that activity in early visual cortex reflects the observer’s interpretation rather than the stimulus itself (e.g., increased V1 activity for perceived faces), suggesting an active, generative process that shapes perception to resolve ambiguity in the pattern. To test whether Generative Inference could replicate this effect, we presented the Rubin stimulus to two versions of a ResNet-50 model: one trained on face recognition and one on object recognition. Despite having identical architectures, the face-trained network inferred face-like contours through feedback, while the object-trained network produced vase-like interpretations. These interpretations did not emerge during the initial feedforward pass, but built up over inference iterations via feedback updates, demonstrating that perception depends jointly on stimulus structure and learned priors.

Another foundational Gestalt principle is figure-ground segregation, which allows the visual system to distinguish objects (figures) from their background. Even low-level texture cues can elicit this effect, as demonstrated in macaque V1 neural recordings where neurons respond more strongly to figure regions than to uniform textures, even when local features are identical—an effect that emerges with a 30-40 ms delay^17;25^, implicating feedback-driven modulation. To test whether Generative Inference could account for this low-level texture-based segregation (in contrast to higher-level shape processing seen in the face-vase illusion), we devised an alternative inference objective called Prior-Guided Drift Diffusion, which requires iteratively moving away from a noisy interpretation of input in intermediate layers (See Supp. B.2). Using Generative Inference by PGDD, we observed a similar enhancement of activations for figure regions in texture-defined stimuli (Figure 4A). The model progressively amplified boundaries and suppressed background texture, recapitulating both the timing and spatial selectivity of figure-ground effects observed in the cortex.

**Figure 4:**
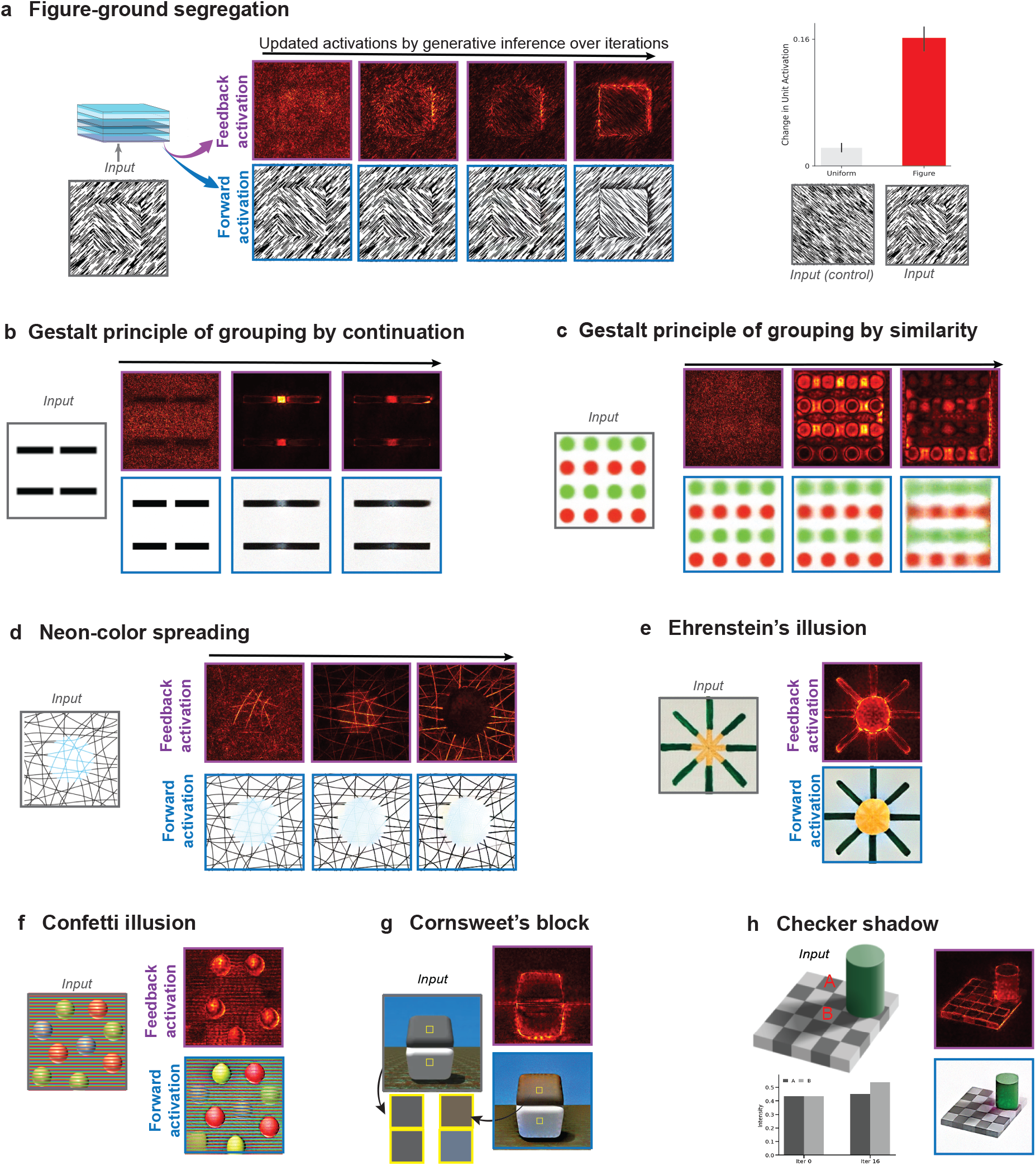
Generative Inference unifies accounts for Gestalt principles of perceptual organization and contextual brightness illusions. **a** Neural responses in animal studies show delayed, enhanced activation in early visual cortex for figure regions relative to uniform texture^17;25^. Generative inference replicates this effect through feedback-driven inference. The top row shows the evolution of activation updates over iterations, revealing progressive enhancement of figure boundaries. The bottom row depicts the corresponding forward activations, demonstrating that feedback refines perceptual interpretations beyond the initial feedforward pass. The bar plot quantifies the increase in unit activation for figure regions compared to uniform backgrounds, aligning with experimental data. **b** (Left) Input stimulus containing intersecting black lines with an embedded blue shape that induces the percept of neon color spreading^36;37^. (Right) Generative inference progressively enhances inferred brightness through feedback-driven updates. **c-f** Generative Inference predicts similar feedback-mediated effects for other classic visual illusions: ^38^ **c** Ehrenstein’s illusion, where viewers typically perceive colored circular regions appearing at the intersection points of the radiating lines. Generative inference demonstrates how these intersections could transform from simple line crossings into filled-in colored circular regions **d** The confetti illusion, where identical colored spheres appear to take on different hues based on the colors of surrounding stripes. Generative inference shows how the model captures this color assimilation effect, enhancing the apparent differences between identically colored objects ^42^ **e** Cornsweet’s brightness contrast, where physically identical regions appear to have different brightness levels ^39;43^, and **f** Checker shadow illusion, where two squares with identical absolute luminance appear different to human observers due to perceived shadows and context^40^. The bar graph quantifies the difference in intensity developed over 16 iterations of generative inference.

We next examined the Gestalt principles of continuation and similarity, which govern perceptual grouping: elements that are near each other or share visual features (e.g., color, shape) tend to be perceived as belonging to the same group^35^. In monkey electrophysiological studies, attentional modulation elicited on one cued element of a grouped set spreads to other uncued elements—suggesting that grouping is implemented via a delayed, feedback-mediated process^28^. Generative Inference replicates this effect: when stimuli contained multiple candidate groupings, feedback-driven updates selectively enhanced activity for elements sharing visual attributes or spatial proximity, even in the absence of explicit labels or attention (Figure 4B-C). These groupings were not hard-coded, but emerged from the network’s learned priors and the structure of the stimulus.

Beyond shape and grouping, visual illusions of brightness, color, and contrast also reflect prior-dependent perception. In neon color spreading, for example, observers report illusory color filling-in across regions that lack physical color^18;36;37^. Similar contextual effects appear in Ehrenstein’s illusion^38^, the Cornsweet effect^39^, and checker shadow illusions^40^, where identical physical stimuli appear different due to their context. Generative Inference accounts for all of these by propagating activation through feedback: color and contrast gradients emerge progressively over inference iterations, driven by the model’s internal priors (Figure 4D-H). While direct electrophysiological recordings for many of these illusions are limited^37^, recent studies in mice have shown delayed V1 responses consistent with neon color filling-in^41^, suggesting that feedback mechanisms similar to those implemented in our framework may underlie these perceptual phenomena in the brain.

Together, these results demonstrate that a single computational mechanism, Generative Inference, can account for a broad class of perceptual phenomena (Figure 1A) traditionally explained by distinct Gestalt principles. This finding challenges the century-old view that perceptual organization requires specialized, hard-wired mechanisms for different visual effects. Rather than invoking separate processes for closure, grouping, or figure-ground segregation, our framework shows that learned priors, integrated via feedback, can dynamically shape perception in a unified and biologically plausible manner.

### Generative Inference accounts for perceptual experiences in absence of sensory inputs

When sensory input is minimal or absent, the brain is still capable of generating rich perceptual experiences. Normally, this manifests as imagination—the ability to internally construct possible percepts or scenarios in the absence of direct stimuli^44;45^. In its pathological form, it may produce hallucinations, where internally generated content is mistaken for sensory input^46^. Both phenomena suggest that perception can be driven entirely by internal priors, without bottom-up input. This raises a key question: can a system trained only on sensory-driven tasks to recognize patterns spontaneously generate coherent structure when no reliable input is available?

To test this, we applied Generative Inference to networks given random noise as input, simulating unstructured sensory input. In the absence of meaningful feedforward activations, we used two alternative inference objectives to update the network’s activations through feedback: the prior-guided drift-diffusion objective, which iteratively removes noise while preserving plausible structure, and the increased confidence objective, which moves representations away from unlikely interpretations while remaining on the learned data manifold, both of which were used above to account for illusions and Gestalt principles. Over successive feedback iterations, updates to activations instructed by generative inference objectives transformed the noise into structured patterns that reflected the network’s learned priors (Fig. 5A-B). Interestingly, the layer at which feedback was applied influenced the granularity of hallucinated structure. Feedback to earlier layers resulted in finer textures and local edges, while feedback to deeper layers yielded more coherent object parts and semantic clusters (Fig. 5B). This mirrors cortical organization in biological vision, where early visual areas encode low-level features and higher areas represent objects and concepts. These results suggest a plausible computational mechanism for how imagined content might be constructed hierarchically in the brain, layer by layer, through iterative integration of priors.

**Figure 5:**
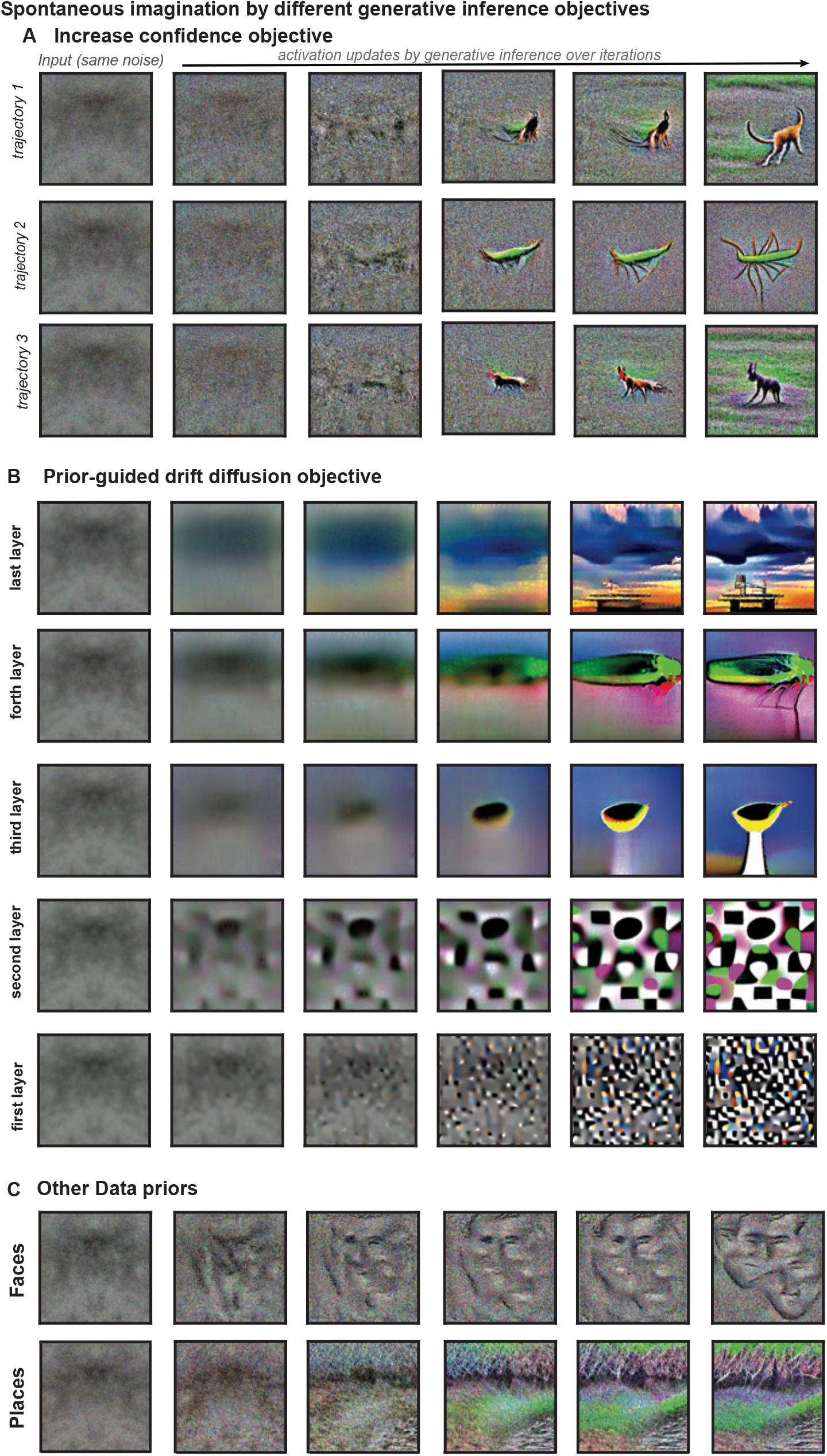
Spontaneous pattern formation by generative inference. When input is random noise, Generative Inference progressively imposes structured patterns reflecting the network’s learned prior distribution. **a** The increase confidence objective drives activation patterns and converges on different patterns depending on the particular trajectory taken on the prior landscape. Note, this objective is necessarily applied at the last layer (one-hot representation of categories) and is back-propagated to all layers. **b** Using prior-driven drift diffusion applied at different layers (and back-propagated to all lower layers) reveals how distinct levels of the visual hierarchy contribute to pattern formation—from basic textures in lower layers to complex objects in higher layers. **c** Networks with identical architectures trained on different datasets (faces, places) generate domain-specific patterns using the same increase confidence objective, demonstrating how priors shape perceptual interpretation in the absence of structured sensory input.

The nature of these hallucinated patterns depended strongly on the network’s training domain (Fig. 5C). A network^23^ trained on object recognition^30^ generated geometric and object-like forms; the same architecture trained on faces^33^ produced fragments resembling facial features; models trained on scenes^34^ generated layouts and spatial configurations. This domain specificity demonstrates that the generative process is not universal or hand-engineered but arises naturally from the network’s learned statistical regularities. In each case, the same feedback pathways used for learning were sufficient to hallucinate new structures, suggesting that imagination, like perception, may result from feedback-driven inference over internal priors.

Taken together, these findings extend the scope of Generative Inference beyond perception into the domain of internally generated experience. Without any architectural change or retraining, standard neural networks exhibit emergent generative capabilities—hallucinating content that reflects their acquired knowledge during training. This supports the idea that imagination and hallucination may arise from the same underlying mechanism as perception: the dynamic reuse of feedback to traverse the learned manifold of likely interpretations, even in the absence of sensory input.

## Discussion

Our findings support a fundamental computational principle: the same feedback pathways that adjust synaptic weights during learning can also dynamically update neural activations during perception. This dual role of feedback explains how visual systems can seamlessly transition between rapid recognition and flexible perceptual inference. When feedback is used to update synaptic weights, it enables learning robust object representations from experience. When repurposed to update activations, it allows integration of learned statistical regularities with incoming sensory evidence, effectively implementing generative inference.

Why does generative inference work in a model that was never explicitly trained to generate? We showed that neural networks optimized to recognize patterns, mapping inputs to labels, do more than just form associations. Implicitly, they internalize the statistical structure of the data distribution. With robustness to noise training or probabilistic inference, this structure becomes accessible through the iterative accumulation of input gradients (see Supplementary A).

Specifically, the network learns to shape its gradients such that updating activations in directions away from unlikely interpretations leads toward regions of higher likelihood under the learned data. In effect, the model’s feedback pathways—typically reserved for error signals—can be repurposed to revise ambiguous or corrupted input in a generative direction, even though no generative objective was ever used.

The computational equivalence we have demonstrated between the gradient of classification loss and the score function of data which is typically used in generative models provides a formal mathematical basis for this unification. By showing that networks optimized for recognition inherently learn statistical regularities that can be accessed through feedback, we bridge traditionally separate paradigms in computational vision and provide a principled explanation for the diverse functions of cortical feedback. While the studied network use backpropagation of error to guide learning (and subsequently inference), which is not considered biologically plausible with our current knowledge about the neural circuits, we suggest that the more general principle is that the signals that guide learning, whether in artificial or in biological networks, will embody priors that can also guide inference.

This insight sets our approach apart from other frameworks that incorporate feedback, such as predictive coding and recurrent neural networks (see Supplementary F). Although these architectures are designed to use feedback connections, end-to-end training does not ensure that feedback carries meaningful prior information. Without an explicit mechanism that aligns feedback with the data distribution, these models often lack the ability to reshape inputs in a generative way. In contrast, our method shows that a standard feedforward network, trained to perform a task, can perform generative inference using its own gradients—offering a simple yet powerful route to modeling perception as a dynamic, prior-guided process.

Generative Inference captures key properties of cortical feedback processing observed across multiple experimental paradigms. At the level of neural circuitry, our framework provides a mechanistic explanation for how internal generative models are implemented in the brain. For decades, psychological and theoretical neuroscience frameworks have posited that the brain must contain internal generative models that enable prediction, inference, and imagination. The Hierarchical Bayesian framework proposed by Lee and Mumford^47^ suggested that feedback plays a key role in combining top-down predictions with bottom-up signals, but direct computational implementations capturing this interaction in neural circuits have remained elusive. Generative Inference reveals that the very same feedback pathways that enable learning can also implement generative capabilities during perception. Our results demonstrate that this integration of priors need not be a separate computation but can be an intrinsic property of cortical processing, providing a concrete computational link between statistical inference and neural mechanisms. This unified principle supports the notion that illusions are not merely perceptual mistakes, but manifestations of the same statistical inference processes that enable robust perception under noise, effective completion of occluded objects, and generative mental imagery in the absence of sensory input.

At the activation level, our model replicates the laminar-specific response patterns observed during illusory contour perception. In both mice and monkeys, neurons in superficial layers and deep layers that receive abundant feedback connections show robust responses to Kanizsa figures, while the primary input layer (L4) remains largely unresponsive to illusory contours. This precise laminar distribution is consistent with our framework’s prediction that feedback-driven inference selectively modulates layers receiving top-down inputs while preserving feedforward sensory representations. Similarly, our model reproduces the characteristic delayed onset of illusory responses (70-100ms in superficial layers versus 30-40ms for real contours), reflecting the iterative accumulation of prior information through feedback loops. Neuroimaging studies have further shown that early visual areas contain decodable information about occluded objects and bistable percepts like the Rubin face-vase illusion, demonstrating that feedback modulates neural representations even in primary sensory areas.

At the perceptual level, our framework accounts for the phenomenological experience of seeing contours, shapes, and complete objects where none physically exist. The feedback-driven activation patterns in our model directly correspond to the reported percepts in human psychophysical experiments. For example, the model generates illusory contours only when inducer segments are properly aligned and within proximity thresholds that match human perceptual boundaries. In the face-vase illusion, the model’s emergent interpretations reflect the bistable percepts reported by human observers, with different priors leading to distinct perceptual outcomes from identical inputs. Through this unified computational principle, our framework reconciles seemingly diverse experimental findings across species and experimental paradigms, suggesting that feedback-driven integration of learned priors may be a fundamental organizing principle of cortical computation in perception.

While our implementation uses backpropagation-trained networks, the theoretical connection between feedback-mediated learning and generative inference holds for any learning algorithm that estimates gradients accurately. This means that biologically plausible alternatives to backpropagation, such as feedback feedforward alignment^48^, local learning rules with eligibility traces, or segregated dendrite models^49^, would support the same computational framework as long as they approximate the true error gradients. Additionally, though our models use continuous-valued units rather than spiking neurons, Poisson spiking neural networks (SNNs) inherently implement the kind of signal averaging required for probabilistic generative inference through their Poisson encoding^50^. This suggests that as SNNs achieve recognition capabilities comparable to conventional deep networks, they could naturally implement generative inference through the same feedback pathways, providing a biologically plausible implementation of smooth inference without requiring explicit averaging over multiple noisy inputs. The stochastic nature of spike timing effectively performs the sampling needed for robust inference in a manner aligned with cortical computation.

Beyond explaining classic perceptual phenomena, our framework points toward a future where artificial systems might approach the fluid intelligence of biological vision—navigating uncertainty, completing fragmented inputs, and synthesizing novel percepts through the same adaptive mechanisms. The marriage of recognition with generative capabilities through feedback suggests a domain-general principle that likely extends beyond vision to audition, touch, and perhaps even abstract reasoning and conceptual thought. As we further explore this computational symmetry between learning and inference, we may ultimately discover that the brain’s ability to both recognize and imagine—to perceive what is and envision what could be—springs from a singular computational principle expressed through the dynamic interplay of feedforward external drive and feedback internal modulation. In this light, Generative Inference offers not just a model of perception but potentially a fundamental organizing principle of intelligence itself.

## 1 Supplementary Materials

The supplementary materials are organized as follows: First, we present the theoretical framework linking pattern recognition to pattern generation, demonstrating how networks trained for classification implicitly learn priors that can be accessed through the learning feedback connections (Section A). We provide mathematical derivations showing the link between classification objective and score-based generative models, and explain how even standard networks encode generative capabilities (C). Next, we detail the Generative Inference algorithm and its variants, explaining how the two main inference objectives—Increased Confidence (B.1) and Prior-Guided Drift-Diffusion (B.2), leverage feedback connections to integrate priors during perception. We also introduce Probabilistic Generative Inference, which enables generative capabilities even in networks without explicit robustness training. The supplementary materials also include details of model architectures, training regimes, and performance metrics (Section E), along with additional analyses that support main figures and additional analyses expanding on the main results. Finally, we provide comparative analyses with alternative frameworks that incorporate feedback processing, including recurrent neural networks and predictive coding networks (Section F).

### A Theory links learning pattern recognition to pattern generation

#### Score-based Generative Models Intuition

Score-based generative models^51^ provide a powerful mathematical framework for modeling complex probability distributions without explicitly parameterizing the entire density function. These models operate by learning the score function—defined as the gradient of the log probability density with respect to the input—which characterizes the local geometry of the data manifold. Rather than directly modeling *p*(*x*), score-based approaches estimate ∇_*x*_ log *p*(*x*), enabling efficient generation through Langevin dynamics or other gradient-based sampling procedures. This formulation offers computational advantages as the score naturally points toward higher-density regions, allowing noise-corrupted samples to be iteratively refined by following the gradient field. The score function effectively serves as a vector field that guides samples toward regions of high probability, making these models particularly effective for tasks requiring navigation of high-dimensional probability landscapes without explicit density evaluation^52;52;53^.

#### Loss Gradients in Classification Networks

In discriminative models like image classifiers, learning proceeds through gradient descent on a loss function that measures prediction error. When a network updates its weights, the gradient of the loss with respect to these weights (∇_*W*_*L*) indicates how connections should change to improve classification performance. This learning signal depends on both the network’s current predictions and the structure of the input data. Critically, calculating these gradients usually requires propagating error signals backward through the network—a process implemented through feedback connections that transmit information from higher to lower layers. These feedback pathways enable credit assignment by distributing the global error to each synapse in proportion to its contribution. In artificial neural networks, this manifests as the backpropagation algorithm or other bio-plausible variants of gradient-based learning, while in biological systems, feedback connections between cortical areas might serve an analogous function^20;21;54^. Thus, feedback plays a fundamental role in learning by carrying the necessary information to guide synaptic modifications, ensuring that each connection updates in a manner that collectively improves the network’s overall performance.

#### Bridging Classification and Generation through Activation Gradients

While the gradient of loss with respect to weights (∇_*W*_*L*) drives learning, the gradient with respect to activations (∇_*h*_*L*) serves a fundamentally different role that has been underexplored, beyond their role in interpretability through attributions^55^. We propose that, under some conditions, these activation gradients in classifiers approximate the score function usually used in generative models. This connection enables the same feedback pathways that facilitate learning to become useful during inference. When a classifier encounters a noisy, corrupted, or unusual input, it can leverage these gradients to navigate toward more probable activation patterns in its representational space—similar to how generative models use score functions to denoise inputs or sample from high-probability regions. By following the gradient of the activation with respect to the loss, the network effectively performs a form of maximum a posteriori inference, refining initial noisy or ambiguous representations toward statistically likely interpretations that align with previously learned patterns. This mechanism provides a principled way for discriminative networks to handle uncertainty through iterative refinement (i.e. generative inference), without requiring a separate generative architecture. Consequently, the feedback connections that initially evolved to enable effective learning may serve a dual purpose: credit assignment during training and perceptual inference during task performance.

The connection between classifiers and generative models has been explored in previous work. Joint Energy-based Models demonstrated that neural networks can simultaneously perform classification and generation by training with both discriminative and generative objectives^56^. Similarly, Santurkar et al.^57^ showed that robust classifiers can be used for image synthesis, revealing unexpected generative capabilities in adversarially trained networks. However, these approaches either require additional generative training objectives during the learning phase or remain largely empirical demonstrations without theoretical foundations. Our work differs in several key respects: First, we provide a theoretical framework explaining why robust classifiers exhibit generative properties through the mathematical connection between input gradients and score functions. Second, we demonstrate that flexible inference objectives are all needed for emergent capabilities, without requiring additional training for generative objectives or architectural modifications.

### B Generative Inference Framework

Generative Inference is a computational principle that enables neural networks to dynamically integrate learned priors with sensory evidence through feedback pathways. The key insight is that the same feedback connections used to propagate error signals during training can be repurposed during inference to guide activations toward regions of higher probability under the network’s learned data distribution. Generative Inference follows a five-step iterative process (Figure **??**A):

1. **Feedforward pass**: Process input through the network to obtain initial activations and class predictions
2. **Inference objective**: Compute the inference objective, for example “increase confidence” or “prior-guided drift diffusion”
3. **Feedback error propagation**: Calculate gradients using the same pathways employed during training, but now with respect to the chosen inference objective rather than the original training objective
4. **Constrained activation update**: Apply gradient-based updates to input or intermediate representations rather than network weights, while constraining changes to remain within an *ϵ*-ball of the original sensory input
5. **Iteration**: Repeat steps 3-4 until convergence or a maximum number of iterations

The critical distinction from standard inference is that feedback connections, typically not used after training, are reactivated to carry gradient information that refines internal representations. This enables the network to move beyond its initial feedforward interpretation and incorporate statistical regularities learned during training.

In principle, any differentiable inference objective can be implemented within this framework. Here, we describe two primary inference objectives that we tested empirically and that are grounded in the theoretical foundation outlined in Section A. The increase confidence objective enhances interpretation certainty by moving away from unlikely classes, while prior-guided drift diffusion recovers structure from corrupted or incomplete inputs. Both objectives leverage the same underlying principle: repurposing learning-derived feedback pathways to implement statistical inference during perception.

#### B.1 Increase confidence objective

The increase confidence objective enhances perceptual certainty by moving representations away from the least likely class interpretation identified during the initial forward pass, while simultaneously preserving consistency with input. The algorithm follows the five-step process illustrated in Figure **??**A:

##### Algorithm 1

Increase Confidence Objective

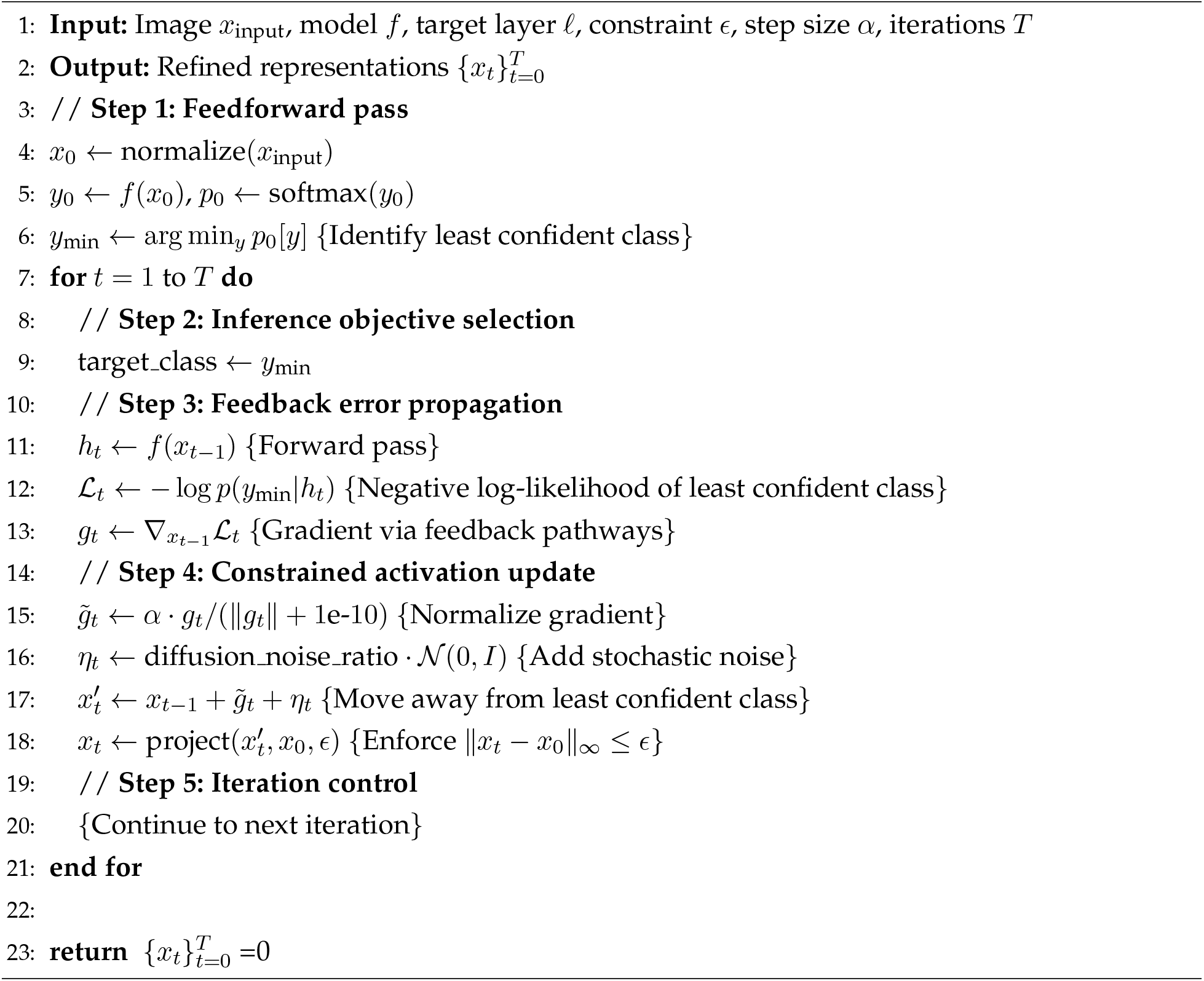

As shown in Figure S1B (left), this objective drives the network away from the least likely class (shown in red among the class probability distribution) toward more confident interpretations. The effectiveness of this approach is demonstrated in Figure S1C, where the increase confidence objective successfully generates illusory contours in Kanizsa figures and produces structured patterns from noise across different object categories.

#### B.2 Prior-guided drift diffusion objective

This objective aims to recover the nearest plausible pattern under learned distribution by iteratively refining representations in latent space. We define the anchor (anti-target) as a noise-corrupted version of the input:

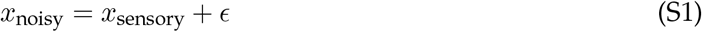

The loss function operates in representation space *r*(·):

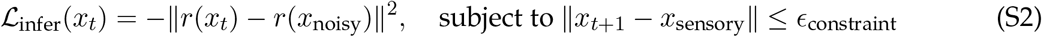

Following the same five-step process (Figure S1A), the algorithm proceeds as:

##### Algorithm 2

Prior-Guided Drift Diffusion Objective

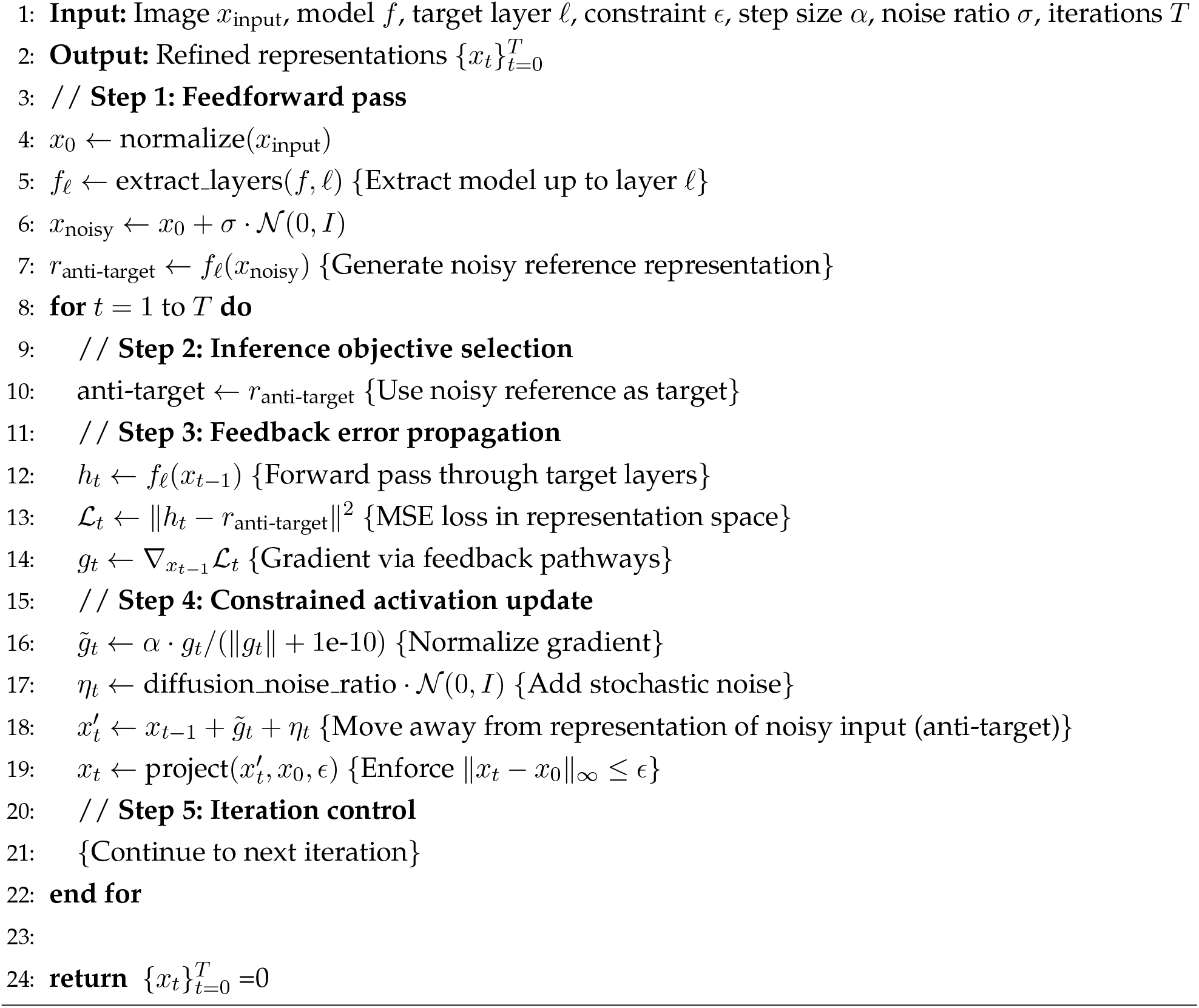

As illustrated in Figure S1B (right), this objective progressively transforms noisy representations into refined, structured patterns. Figure S1D demonstrates the results across different iterations, showing how the algorithm gradually imposes the network’s learned statistical regularities to generate coherent visual patterns from corrupted inputs. Both objectives leverage the same feedback pathways established during training, repurposing them to guide activations toward regions of higher probability under the network’s learned prior distribution.

**Figure S1:**
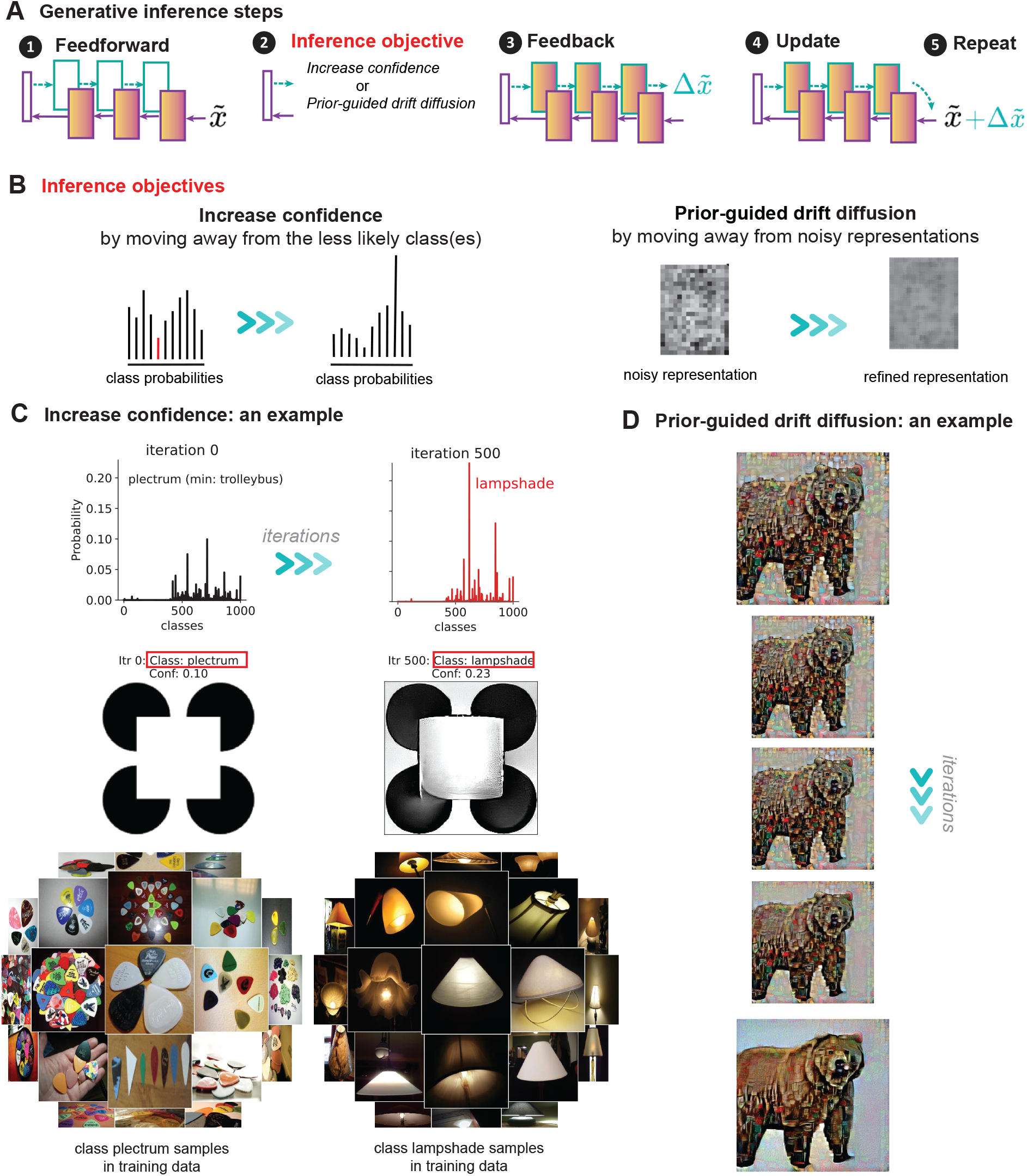
Generative Inference Objectives. **A** The five-step generative inference algorithm: (1) feedforward pass, (2) objective selection, (3) feedback gradient computation, (4) activation update, and (5) iteration until convergence. **B** Comparison of the two inference objectives: increase confidence (left) moves away from least likely classes identified in the initial forward pass, while prior-guided drift diffusion (right) refines noisy representations toward clean patterns. **C** Results from increase confidence objective showing successful generation of illusory contours (Kanizsa figures) and structured patterns from noise across different training categories. **D** Results from prior-guided drift diffusion objective demonstrating progressive refinement of corrupted inputs over multiple iterations, revealing the emergence of coherent visual patterns through feedback-driven prior integration.

### C Probabilistic generative inferene

As established in our theoretical framework (Section C), neural networks do not necessarily require adversarial robustness training to enable generative capabilities through feedback. Even networks trained with standard procedures implicitly encode the score function of the data distribution, albeit with a smaller stability radius compared to adversarially trained networks.

#### Accessing the Score Function in Standard Networks

The idea here is that by averaging gradients over multiple noise-corrupted versions of an input, we can approximate the score function even in standard neural networks:

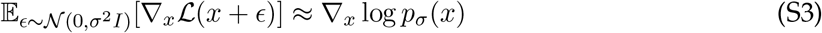

Where *p*_*σ*_(*x*) represents the data distribution convolved with Gaussian noise. This approach allows us to access the generative capabilities inherent in any neural network without requiring explicit robustness training. The averaging effectively cancels out the class-dependent term in the gradient decomposition, isolating the data prior term that guides generative processes.

To validate this approach, we implemented probabilistic generative inference across several network architectures with standard training (no adversarial robustness). At each iteration, we:

1. Generate multiple noise-corrupted versions of the current state
2. Compute gradients for each noisy sample
3. Average these gradients to approximate the score function
4. Update the activation patterns based on this average gradient

Figure S4 demonstrates that standard neural networks (Convolutional: ResNet50, Transformer: vit-l-16) without adversarial training generate illusory contours and figure-ground modulation when using probabilistic generative inference. The quality of inference improves with the number of noise samples used for gradient averaging, with diminishing returns beyond approximately 10-15 samples.

**Figure S2:**
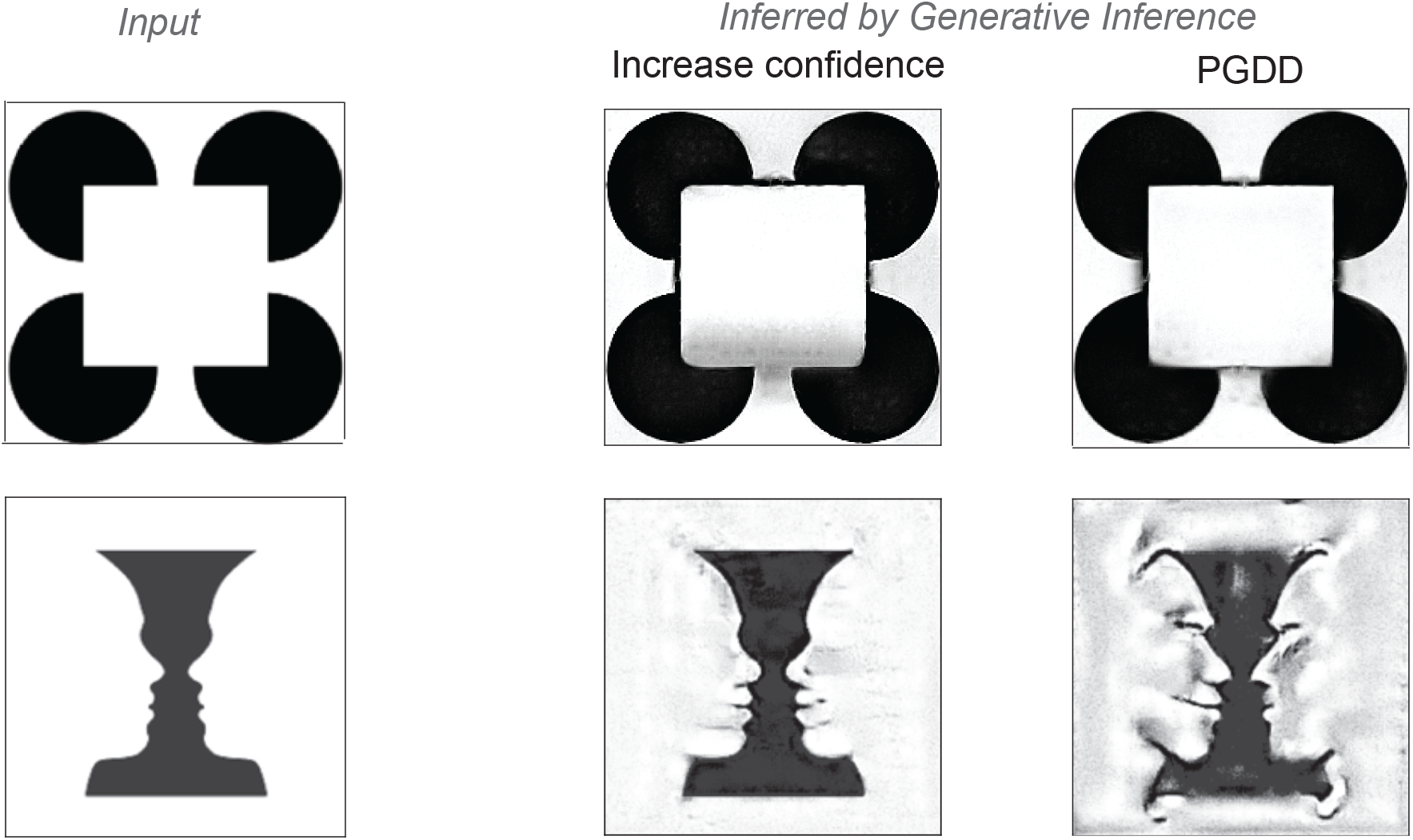
Increase confidence and PGDD algorithms arrive at different variants of acceptable account for perceptual experiences, here Kanizsa square and Rubin’s vase shown.

Interestingly, the gradient averaging procedure required by probabilistic generative inference aligns naturally with properties of biological neural networks. Spiking neural networks with Poisson-distributed action potentials inherently implement a form of noise sampling and averaging through their stochastic firing patterns.

The variance in spike timing effectively performs implicit sampling from a distribution of inputs, analogous to our explicit averaging over noise-corrupted inputs. This provides a biologically plausible mechanism for implementing probabilistic generative inference without requiring multiple explicit forward passes. As suggested in recent computational neuroscience research^50^, the brain’s intrinsic noise may actually serve a computational purpose by enabling this implicit sampling process necessary for generative inference.

While probabilistic generative inference works for standard networks, our experiments confirm that adversarially trained networks still provide superior results with fewer iterations. This is because:

1. The stability radius induced by standard training is smaller than in robust models
2. The local Gaussian approximation holds more reliably in adversarially trained networks
3. Robust networks require fewer noise samples to achieve stable inference

Nevertheless, the emergence of generative capabilities across both training paradigms supports our central theoretical claim: the dual role of feedback for both learning and inference is a fundamental principle that transcends specific training methodologies.

The effectiveness of probabilistic generative inference across different network architectures and training regimes suggests that generative capabilities may be a universal property of networks trained for visual recognition. This broadens the applicability of our framework beyond models specifically trained for robustness, potentially making generative inference accessible as a general-purpose mechanism for enhancing perception and explaining illusions across a wide range of existing vision models.

This finding also strengthens the biological plausibility of our approach, as it suggests that the brain’s generative capabilities need not depend on specialized robustness mechanisms but could emerge naturally from the statistical learning processes inherent in sensory systems.

### D Theoretical Foundation of Generative Inference

#### D.1 Gradient Decomposition

For a classifier parameterized by *θ*, the loss is typically the negative log-likelihood:

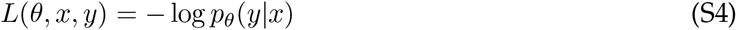

Using Bayes’ rule with a uniform prior *p*(*y*), we write:

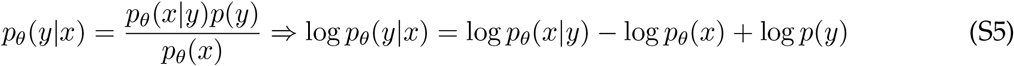

Taking the gradient with respect to *x*:

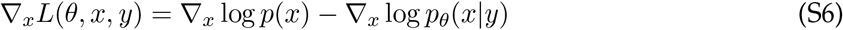

This decomposition reveals that input gradients contain two components: the **data distribution score** ∇_*x*_ log *p*(*x*) and the **conditional class score** ∇_*x*_ log *p*_*θ*_(*x*|*y*). This relationship holds for any class *y*, providing the mathematical foundation for generative inference.

#### D.2 Flatness of Conditional Score around least-likely classes

The key insight enabling generative inference by increasing confidence is that the conditional score for the least likely class becomes locally flat. This flatness can be achieved through two mechanisms:

##### Robust Training

Networks trained with gradient regularization (e.g., adversarial training) explicitly penalize large input gradients, creating smooth, flat regions around training examples. For correctly classified inputs, the least likely class represents a distribution the input demonstrably does not belong to, resulting in locally constant (flat) conditional probability and thus vanishing gradients:

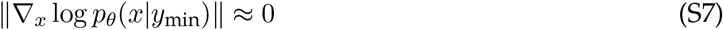

##### Probabilistic Inference

Even in standard networks, averaging gradients over noise-corrupted inputs approximates the score function while smoothing out sharp conditional variations. This averaging effectively creates the same flatness property:

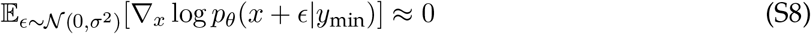

#### D.3 Generative inference by Increase Confidence effectively runs gradient ascent on the learned data distribution

Given conditional flatness for the least likely class, the gradient decomposition from Equation 6 simplifies to:

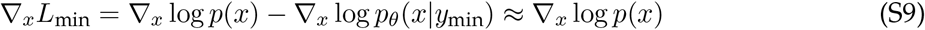

**Figure S3:**
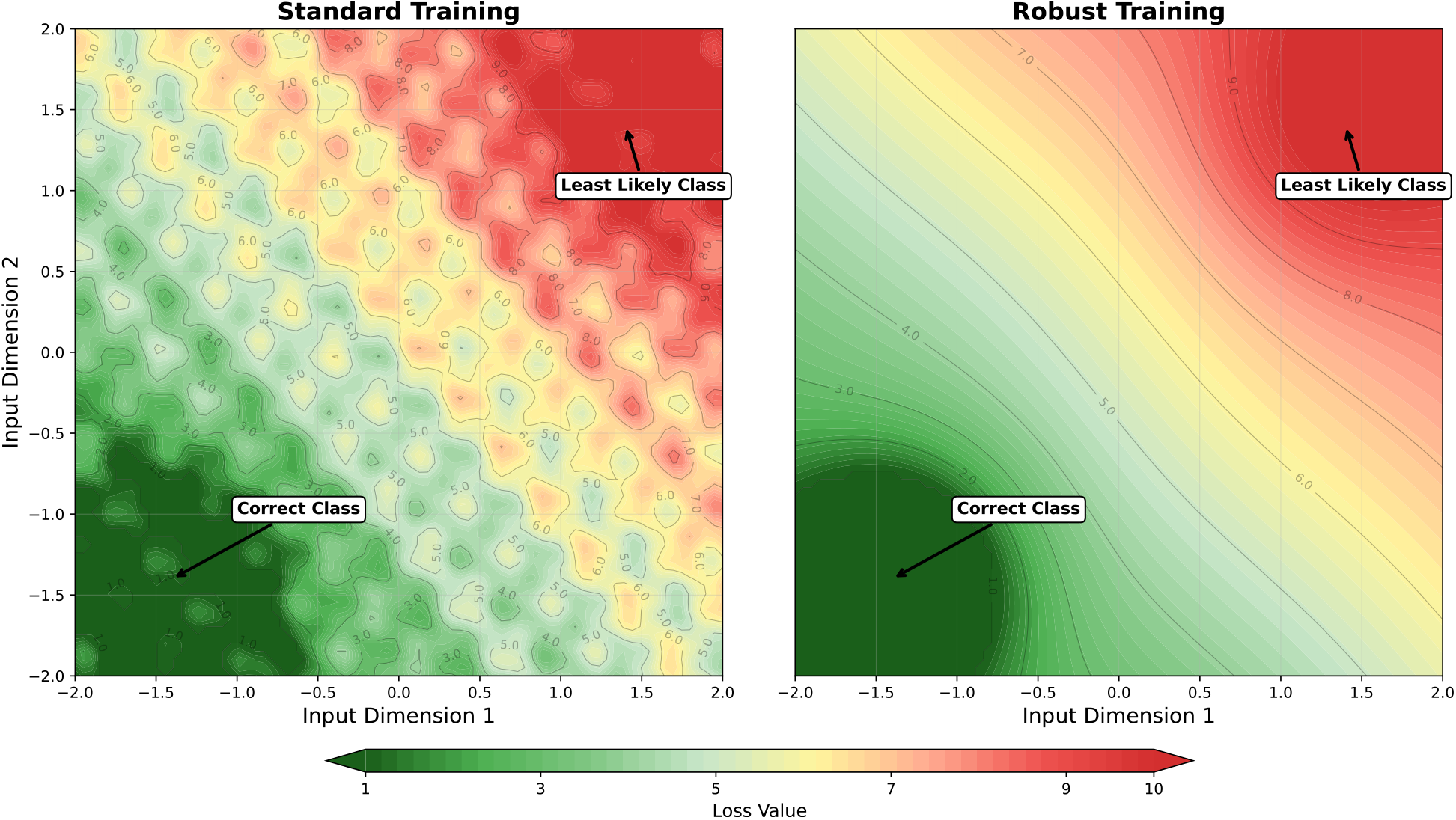
Schematic illustration of loss landscapes demonstrating conditional score flatness. Conceptual visualization showing how different training paradigms affect the geometric properties of loss landscapes in input space. **Left:** Standard training produces spiky, irregular loss landscapes with steep gradients and sharp transitions between class regions. The conditional probability for the least likely class varies across input space, requiring probabilistic inference (smoothing) to ensure access to data score. **Right:** Robust training creates smooth, regular contours with large flat regions around the least likely class. This schematic illustrates the theoretical principle that robust training ensures ∥∇_*x*_ log *p*_*θ*_(*x* | *y*_min_) ∥ ≈ 0 (Equation S7), allowing the gradient decomposition to simplify to ∇_*x*_*L*_min_ ≈ ∇_*x*_ log *p*(*x*) (Equation S9). The smooth landscape geometry enables reliable gradient ascent on the learned data distribution through the increase confidence objective.

Therefore, the increase confidence update:

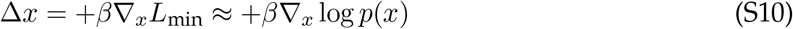

effectively performs **gradient ascent on the learned data distribution**. Moving away from the least likely interpretation directly increases probability under the data prior.

#### D.4 Generative inference by Prior-Guided Drift Diffusion

The Prior-Guided Drift Diffusion (PGDD) objective draws direct inspiration from denoising score matching and Denoising Diffusion Probabilistic Models (DDPMs)^58^. This connection reveals a fundamental parallel between diffusion-based generation and the implicit capabilities of robust classifiers^24;29^.

In DDPMs, networks learn to predict added noise by minimizing:

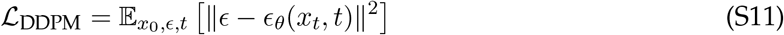

where 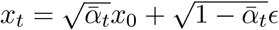. This noise prediction objective implicitly learns the score function of the data distribution, enabling iterative generation through gradient-based sampling.

##### PGDD Objective

We propose that robust classifiers can be used for similar iterative refinement. Given a corrupted input *x*_noisy_ = *x*_clean_ + *E*, PGDD maximizes the MSE loss between current input representations and the noisy reference, effectively moving away from a noisy representation of the current input while staying close to the original input:

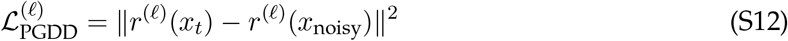

The update rule is:

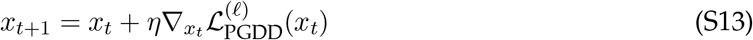

##### Theoretical Foundation for PGDD

To understand why PGDD enables generative inference, we first derive the gradient of inference loss w.r.t input as was used during PGDD and then we show that how it leverages the implicit denoiser structure to remove the noise in each iteration and become closer to data manifold.

##### Gradient of loss w.r.t input in PGDD

For ***ϵ***∼ *𝒩* (0, *σ*^2^**I**) and layer *r*(·),

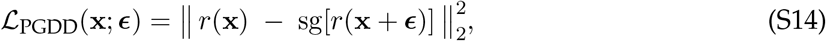

where sg[·] stops gradients through its argument. Let **J**_*r*_(**x**)= ∇_**x**_*r*(**x**). Then

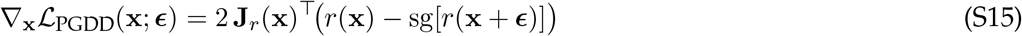

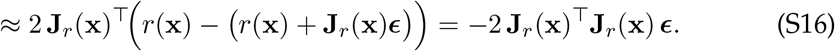

Since the goal is to move away from the noisy representation, we ascend on (S14):

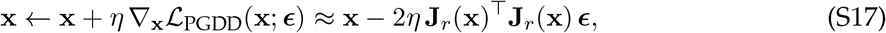

which applies the denoising step 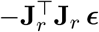. Thus, **J**^T^**J** emerges naturally from the simple objective of moving away from noisy representations.

##### Implicit denoiser in robust classifiers

A growing body of work suggests a tight connection between the spectral structure of network Jacobians and adversarial robustness. Even in standard training, neural networks tend to exhibit low–effective–rank Jacobians: empirical studies have shown that only a handful of singular values carry most of the sensitivity, while the majority are near zero^59;60^. Robust training appears to sharpen this spectral decay.

Hoffman et al.^61^ further showed that robustness can be explicitly encouraged through Jacobian regularization, directly suppressing small singular directions. Most directly, Nassar et al.^62^ enforced a 1*/n* eigenspectrum in hidden-layer covariance matrices, motivated by observations in biological V1, and found that such spectral shaping significantly improves adversarial robustness. Together, these findings support the interpretation that adversarial training yields low–rank Jacobians *J*, concentrating sensitivity into a small number of dominant directions. Since *J*^*T*^*J* inherits the same spectrum, it exhibits corresponding low–rank structure: the operator preserves signal-aligned components while strongly contracting orthogonal noise. This explains why *J*^*T*^*J* acts as an implicit denoiser in robust networks, filtering perturbations in irrelevant directions while retaining task-relevant structure.

Thus, just as DDPMs enable generation through learned noise estimation, PGDD enables robust classifiers to generate or complete patterns through implicit score function estimation encoded in their gradient structure. Both increase confidence, and PGDD performs iterative refinement guided by learned statistical regularities, demonstrating that the boundary between discriminative and generative modeling is more fluid than traditionally assumed.

### E Models and algorithms

#### Architectures and weight optimizations

In this study, we used various models under two training regimes: standard training and robust to input noise. Unless otherwise stated the default model we used was a ResNet50 trained to be robust to adversarial noise in ImageNet 1000-way classification. However, we used other architectures such as (VGG16, ResNet18, WideResNets, etc) other datasets (VGGFace2 and Places365). Below is a table summarizing the training parameters and performances for all the models used in this study.

**Figure S4:**
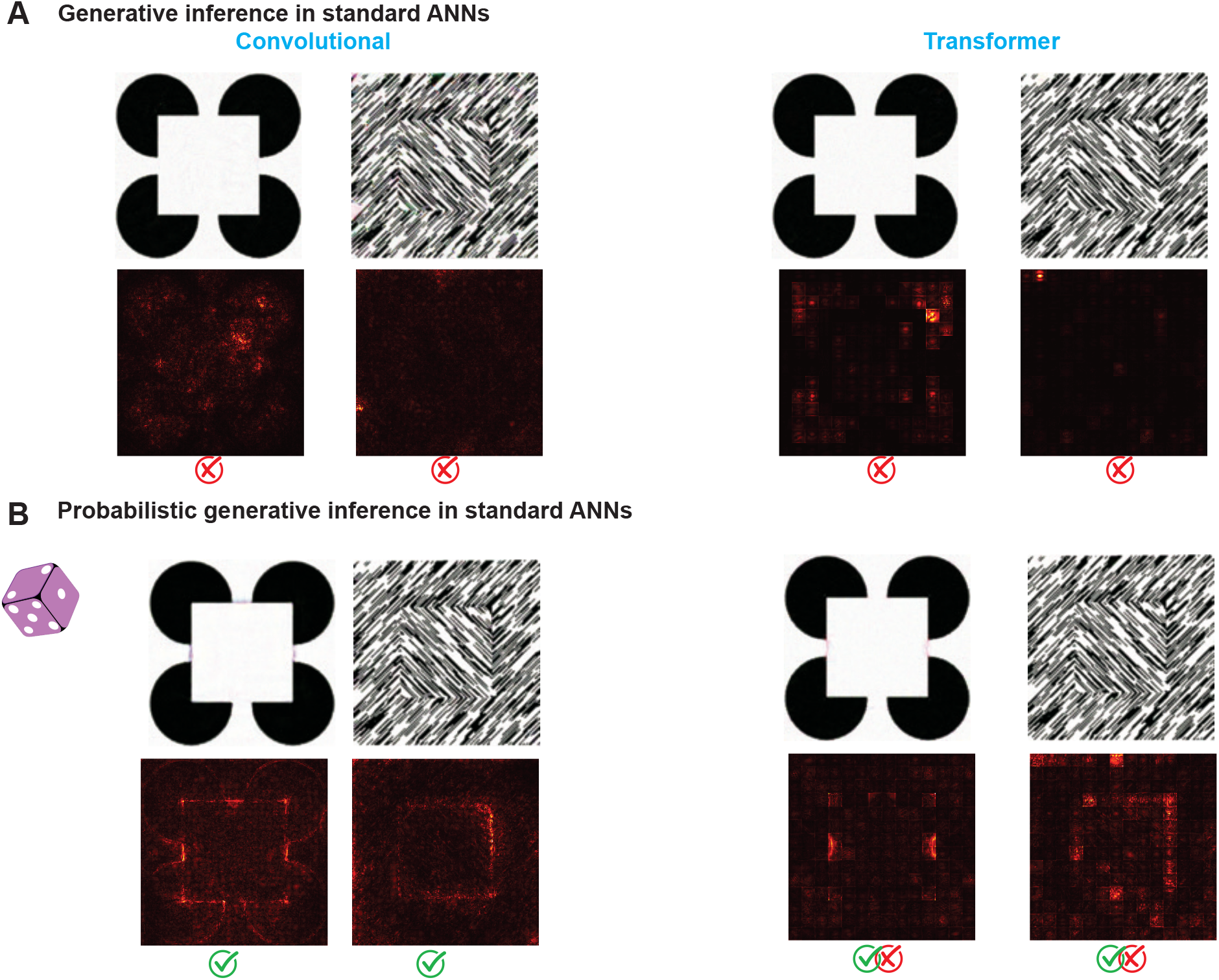
Probabilistic generative inference. **A** Standard neural networks (convolutional and transformer architectures) without adversarial training fail to generate illusory contours and figure-ground modulation when using basic generative inference, as indicated by the absence of the expected activation patterns (compare with Figure 2 and Figure **??**). **B** However, when implementing probabilistic generative inference explained in C, these same networks generate illusory contours and figure-ground modulation, although less evident in the transformer-based model. Convolutional model: ResNet50, and Transfomer model: ViT-Large-patch16 both trained on imagenet classification without any adversarial training.

**Table S1:** Model architectures and training configurations used in the study. The table shows validation accuracy for different neural network architectures trained on various datasets with different levels of adversarial robustness. The *ϵ* parameter indicates the strength of adversarial training (*ϵ* = 0.00 corresponds to standard training without robustness). Chance-level performance is 0.1% for ImageNet (1000 classes), 0.2% for VGGFace2 (500 classes), and 0.28% for Places365 (365 classes). The starred ResNet-50 model (*ϵ* = 3.00, ImageNet) represents the primary model used throughout most experiments unless otherwise specified. Robust training generally reduces standard accuracy but enables the gradient properties necessary for generative inference without resorting to probabilistic inference necessary for running generative inference in non-robust networks.

| Architecture | Dataset | $\epsilon$ | Robust | (Robust) Accuracy (%) |
| --- | --- | --- | --- | --- |
| ResNet-50 * | ImageNet | 3.00 | Yes | 38.00 |
| ResNet-50 | ImageNet | 0.00 | No | 76.65 |
| ResNet-50 | ImageNet | 1.00 | Yes | 57.86 |
| ResNet-50 | ImageNet | 2.00 | Yes | 47.90 |
| ResNet-50 | ImageNet | 5.00 | Yes | 28.70 |
| ResNet-18 | ImageNet | 0.00 | No | 69.79 |
| ResNet-18 | ImageNet | 3.00 | Yes | 42 |
| wide-ResNet50-2 | ImageNet | 0.00 | No | 78.50 |
| wide-ResNet50-2 | ImageNet | 3.00 | Yes | 54 |
| vgg16-bn | ImageNet | 0.00 | No | 71.6 |
| vgg16-bn | ImageNet | 3.00 | Yes | 40 |
| vit-large-patch16 | ImageNet | 0.00 | No | 77 |
| ResNet-50 | VGGFace2 | 0.50 | Yes | 77.73 |
| ResNet-50 | Places365 | 3.00 | Yes | 35.15 |

**Table S2:** Generative Inference configuration as shown in main figures. This table details the specific parameters used to produce the visual results shown in the main paper. For each figure, we specify: the input stimulus type, inference algorithm (Increase confidence or PGDD : Prior-Guided Drift-Diffusion), loss function (MSE : Mean Squared Error), network layer used for loss calculation, noise parameters (noise ratio and diffusion noise ratio), constraint threshold (eps), and update step size.

| Figure | input | loss_infer | loss_function | loss_layer | drift_noise | diffusion_noise | eps | update_rate |
| --- | --- | --- | --- | --- | --- | --- | --- | --- |
| Figure 2 | Kanizsa | IncreaseConfidence | CE | last layer | 0.01 | 0.003 | 5 | 0.5 |
| Figure 3 | Figure-ground | PGDD | MSE | layer3 | 0.5 | 0.005 | 20 | 0.8 |
| Figure 3 | Uniform BG | PGDD | MSE | layer3 | 0.5 | 0.005 | 20 | 0.8 |
| Figure 3 | Neon color | PGDD | MSE | layer3 | 0.5 | 0.003 | 3 | 1 |
| Figure 3 | Ehrenstein | PGDD | MSE | layer3 | 0.5 | 0.005 | 20 | 0.8 |
| Figure 3 | Confetti illusion | PGDD | MSE | layer3 | 0.7 | 0.01 | 0.5 | 1 |
| Figure 3 | Cornsweet's Block | PGDD | MSE | layer3 | 0.5 | 0.005 | 20 | 0.8 |
| Figure 3 | Checker shadow | PGDD | MSE | last layer | 0.1 | 0.005 | 0.3 | 1 |
| Figure 3 | Grouping by Similarity | PGDD | MSE | layer3 | 0.1 | 0.005 | 4 | 1 |
| Figure 3 | Grouping by Continuity | PGDD | MSE | layer3 | 0.1 | 0.005 | 4 | 0.4 |
| Figure 4 | Face-vase (object prior) | PGDD | MSE | last layer | 0.7 | 0.005 | 4 | 1 |
| Figure 4 | Face-vase (face prior) | PGDD | MSE | layer4 | 0.51 | 0.01 | 20 | 0.5 |
| Figure 5 | Imagination (A) | PGDD | MSE | last layer | 0.5 | 0.003 | 40 | 5 |

**Figure S5:**
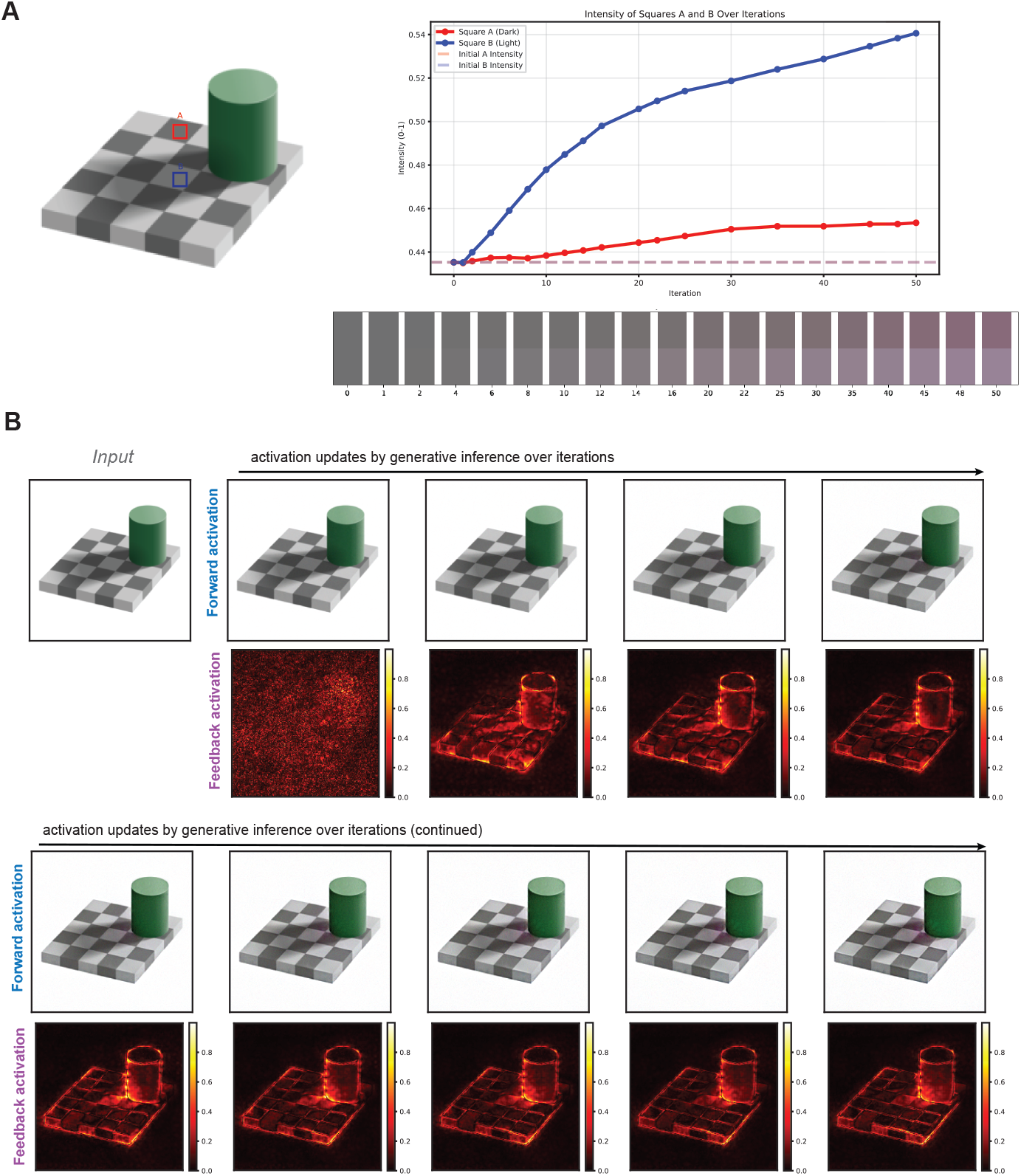
Generative Inference captures the checker shadow illusion through feedback-mediated processing. Generative Inference replicates the classic checker shadow illusion where squares A and B appear to have different luminance despite being physically identical. **A** Progressive changes in the updated forward activation intensity difference between the two squares over 50 iterations of generative inference, demonstrating how the network develops distinct representations that align with human perceptual experience. **B** Evolution of activation patterns showing how feedback pathways iteratively refine the network’s interpretation. Forward activation (top row) shows the updated activation after applying the feedback adjustment in each iteration to increase confidence, while feedback activation (bottom row) shows the feedback (gradient of inference objective) at each iteration. The network progressively amplifies the difference in forward activation between identical squares based on contextual shadow cues, replicating the illusory perception reported by human observers.

**Figure S6:**
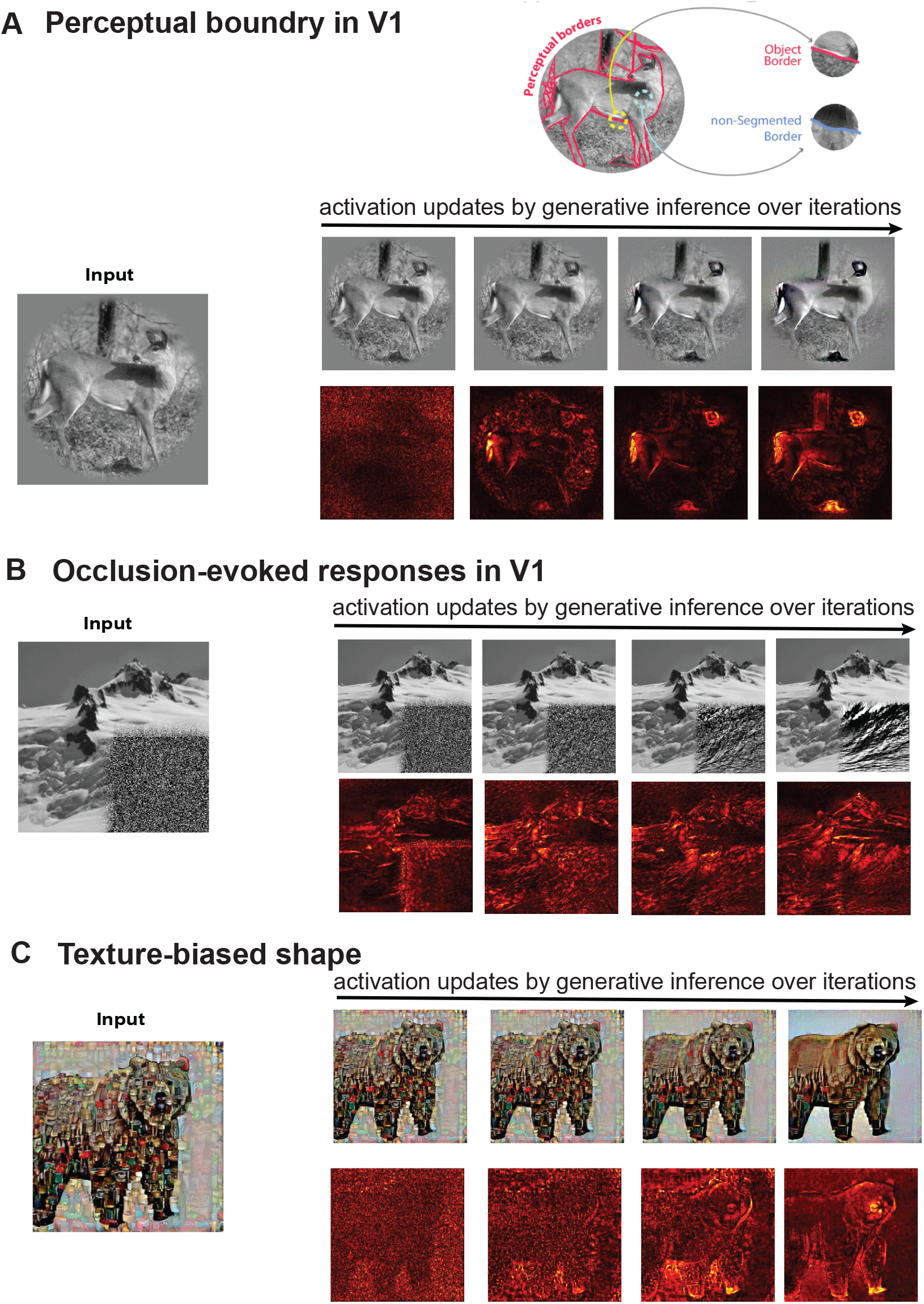
Generative Inference extends beyond illusions. **A** Encoding of perceptual boundary in primary visual cortex. When viewing natural images with complex object boundaries, V1 neurons show enhanced responses to perceptually relevant borders that separate objects from backgrounds, even when local contrast and orientation features are controlled^63^. Generative Inference replicates this boundary enhancement through feedback-driven updates, progressively strengthening object contours (red activation maps) while maintaining naturalistic image structure across iterations. **B** Completion of occluded image regions. V1 neurons respond selectively to occluded portions of natural scenes, encoding information about the hidden content through contextual influences^10–12^. Our model demonstrates similar completion behavior, inferring plausible structure in occluded regions through iterative feedback updates that integrate surrounding visual context with learned priors. **C** Shape completion under texture-shape conflicts. When shape and texture cues conflict, as demonstrated in studies showing ImageNet-trained networks exhibit strong texture bias^64^, Generative Inference can overcome this bias through feedback processing. The model progressively enhances shape-based representations (evident in both forward activations and feedback patterns) while suppressing irrelevant texture information.

### F Other accounts for recurrence fail to capture neural signatures of generative inference: recurrent neural networks and predictive coding

We evaluated Generative Inference alongside two leading models that incorporate feedback processing for visual recognition. CORnet-S offers a biologically-plausible architecture that balances simplicity with performance, using recurrent connections to implement iterative processing while maintaining relatively few parameters compared to state-of-the-art vision models. Its design specifically aims to capture key organizational principles of the primate visual cortex. Meanwhile, PCN implements a fundamentally different computational principle based on predictive coding theory, where adjacent layers interact through bidirectional connections, feedback connections transmit predictions from higher to lower layers, while feedforward connections carry prediction errors that reflect the mismatch between predictions and actual representations. These models serve as strong comparative benchmarks as both were trained on ImageNet for object recognition and have demonstrated competitive performance despite their distinct architectural approaches.

A critical distinction between these frameworks lies in how feedback is used during training versus inference. All three models (including ours) use backpropagation during training, where feedback pathways transmit error signals for weight updates. However, during inference, these pathways serve fundamentally different purposes:

In CORnet-S, recurrent connections operate with fixed weights during inference and don’t explicitly utilize the same feedback pathways that carried error signals during training. Instead, they implement repeated operations within cortical areas, effectively extending processing depth. In PCN, feedback connections during inference carry top-down predictions between adjacent layers, creating a specialized mechanism for prediction error minimization rather than directly leveraging the gradient information used during training. In contrast, Generative Inference makes a direct connection between learning and inference by repurposing the exact same feedback pathways that carried error signals during training. During inference, these pathways now transmit gradients that guide activations toward regions of higher probability under the learned pior.

**Figure S7:**
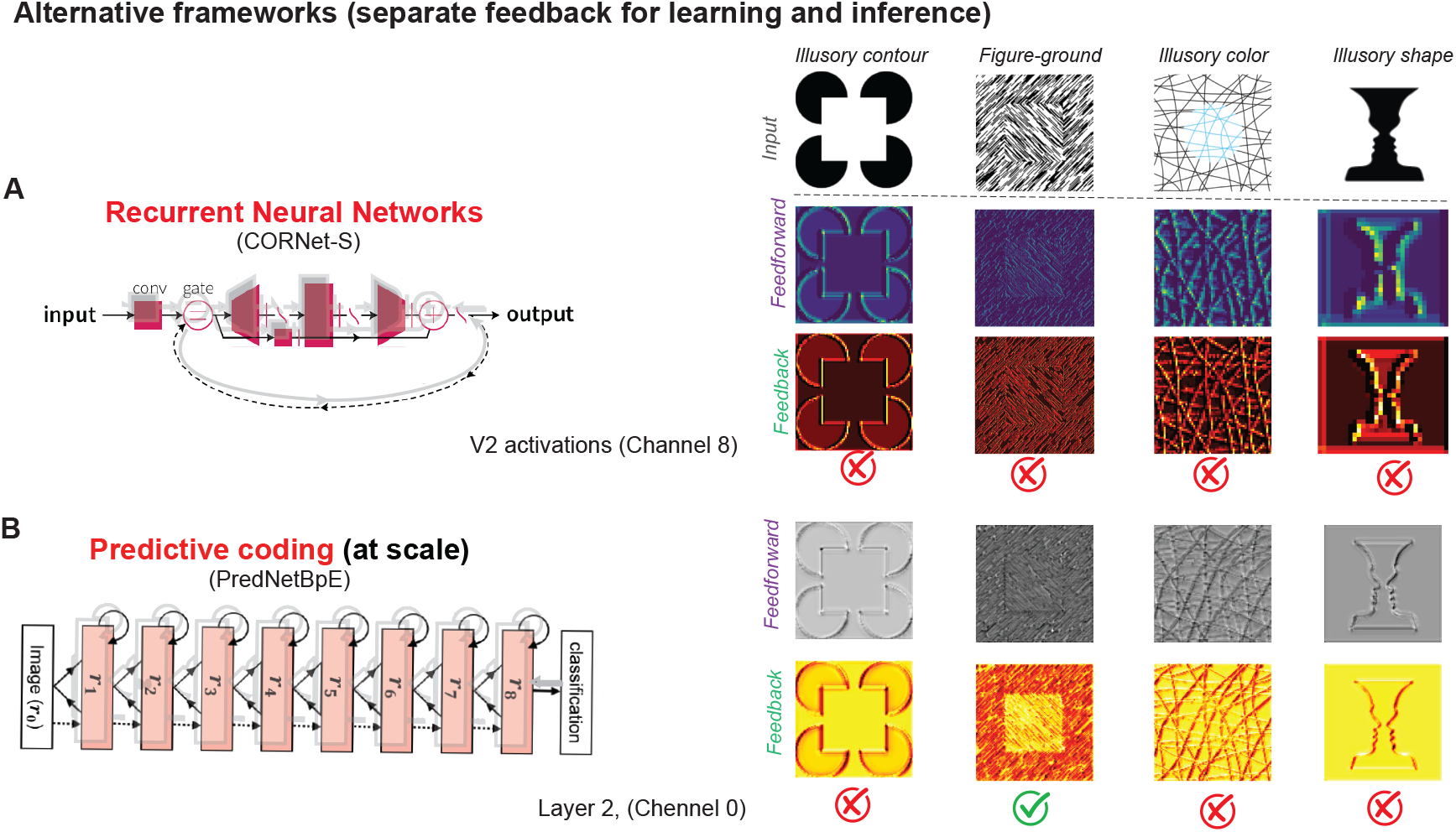
Alternative frameworks implementing separate feedback mechanisms for learning and inference fail to capture the full range of perceptual phenomena. Instances of **A** recurrent neural networks (top) such as CORnet-S^65^ show minimal illusory processing despite strong recurrence, while **B** Predictive Coding Networks such as PCN (bottom)^66^, capture only figure-ground segregation but fail on other instances of neural signatures of prior integration.

